# Unlocking Expanded Flagellin Perception Through Rational Receptor Engineering

**DOI:** 10.1101/2024.09.09.612155

**Authors:** Tianrun Li, Esteban Jarquin Bolaños, Danielle M. Stevens, Hanxu Sha, Daniil M. Prigozhin, Gitta Coaker

## Abstract

The surface-localized receptor kinase FLS2 detects the flg22 epitope from bacterial flagella. FLS2 is conserved across land plants, but bacterial pathogens exhibit polymorphic flg22 epitopes. Most FLS2 homologs possess narrow perception ranges, but four with expanded perception ranges have been identified. Using diversity analyses, AlphaFold-Multimer modeling, and amino acid properties, key residues enabling expanded recognition were mapped to FLS2’s concave surface, interacting with the co-receptor and polymorphic flg22 residues. Synthetic biology enabled engineering of expanded recognition from QvFLS2 (*Quercus variabilis*) and FLS2^XL^ (*Vitis riparia*) into homologs with canonical perception. Evolutionary analyses across three plant orders showed residues under positive selection aligning with those binding the co-receptor and flg22’s C-terminus, suggesting more alleles with expanded perception exist. Our experimental data enabled the identification of specific receptor amino acid properties and AlphaFold3 metrics that facilitate predicting FLS2-flg22 recognition. This study provides a framework for rational receptor engineering to enhance pathogen restriction.

## Introduction

Pattern Recognition Receptors (PRRs) on the plant cell surface can detect pathogens as non-self and mount a defense response. These receptors can recognize conserved microbial features known as microbe-associated molecular patterns (MAMPs)^1,2^. Leucine-rich repeat receptor kinases (LRR-RKs) are the most abundant and extensively studied type of surface receptor^2,3^. LRR-RKs have the capability to directly bind proteinaceous MAMPs^4^. Upon MAMP recognition, plant immune responses are initiated, including calcium influx, reactive oxygen species (ROS) production, phosphorylation of mitogen-activated protein kinase (MAPK) cascades, transcriptional reprograming and callose deposition^5^.

One of the most well-studied bacterial-sensing PRRs is the LRR-RK Flagellin-sensing 2 (FLS2)^6^. FLS2 recognizes a conserved 22-amino acid immunogenic epitope (flg22) from the bacterial flagellin protein monomer^6,7^. Upon flg22 recognition, the SERK (Somatic Embryogenesis Receptor Kinase) family co-receptor BAK1 binds to the C-terminus of flg22, forming a complex with FLS2^8^. FLS2 is present across most land plants and has been characterized in diverse species, but most characterized homologs recognize epitopes similar to the canonical flg22 found in *Pseudomonas aeruginosa* (Pae)^9–12^. However, many α-and some β-proteobacteria possess non-immunogenic flg22 variants that escape immune recognition^13,14^.

Recently, FLS2 homologs with expanded ligand specificity have been discovered. For instance, FLS2 from a wild grapevine species *Vitis riparia* (FLS2^XL^)^15^, star jasmine *Trachelospermum jasminoides* (TjFLS2)^16^, and Chinese cork oak *Quercus variabilis* (QvFLS2)^16^ recognize polymorphic flg22 epitopes from *Agrobacterium tumefaciens*. Similarly, FLS2 homologs from *Glycine max* (GmFLS2s) specifically recognize flg22 from *Ralstonia solanacearum,* a devastating soil-borne bacterial pathogen^17^. Transgenic expression of FLS2^XL^ and the GmFLS2 receptor complex conferred resistance against *Agrobacterium* and *Ralstonia* pathogens in tobacco and tomato^15,17^, respectively, demonstrating the potential of leveraging expanded flg22 specificity for disease control.

In this study, we analyzed the recognition profile of FLS2 homologs against polymorphic flg22 epitopes. Three novel FLS2 homologs (FLS2^XL^, TjFLS2 and QvFLS2) exhibit distinct recognition profiles and response magnitudes. Using a combination of sequence conservation analysis and AlphaFold-Multimer modeling, we developed an approach to harness polymorphic residues on the receptor binding interface to engineer expanded ligand specificity. Evolutionary analysis and refined assessment of amino acid properties bolstered the engineering approach, demonstrating that certain FLS2 residues interacting with polymorphic flg22 residues and co-receptors are under positive selection. This indicates an evolutionary adaptation of FLS2 towards expanded ligand specificity and outlines a rational approach for receptor engineering.

## Results

### Natural FLS2 homologs exhibit unique but expanded flg22 ligand specificity

To explore the extent of expanded ligand perception in different FLS2 homologs, we selected diverse flg22 epitopes from a comparative genomics study of 4,228 plant-associated bacterial genomes^18^. Ten polymorphic flg22 variants from important plant pathogens, representing α, β, and γ-proteobacteria were chosen (Fig. 1a-b, Extended Data Fig. 1). These epitope variants exhibited 31.8 to 68.2% amino acid similarity to the consensus flg22 epitope from *P. aeruginosa* (Pae) and harbored C-terminal polymorphisms (Fig. 1c, Extended Data Fig. 1a). The selected flg22 variants exhibited diverse levels of abundance, but each was found in multiple bacterial species (Fig. 1b, Extended Data Fig. 1c). Although most variants are present in multiple species, we used the most prevalent pathogenic species to represent each flg22 variant. (Extended Data Fig. 1b).

**Figure 1.**
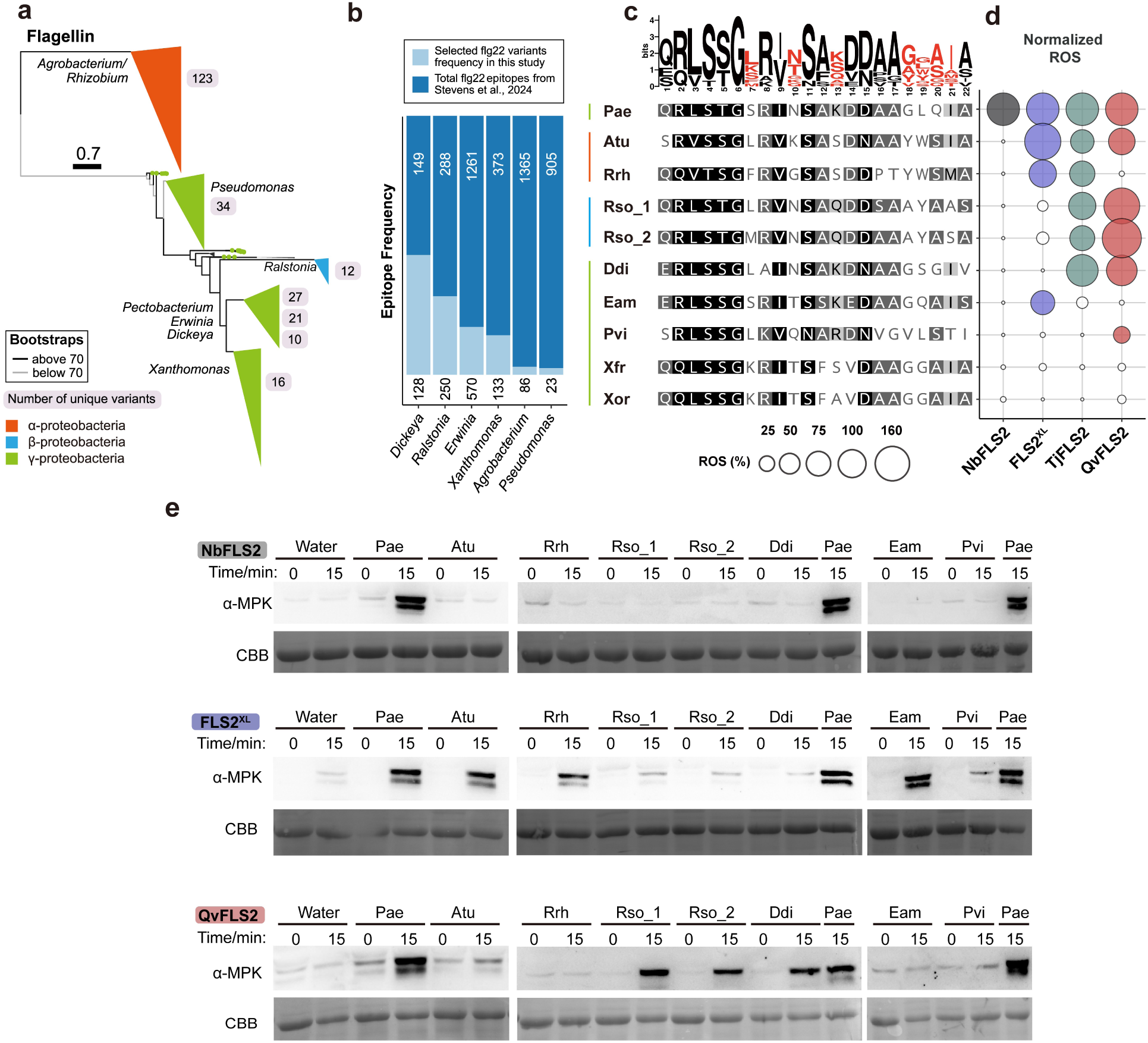
FLS2 homologs exhibit distinct perception range and magnitude. **a,** Maximum-likelihood, midpoint-rooted phylogenetic tree of bacterial flagellin. Major clades are collapsed in a genera-dependent manner and colored based on taxonomic classes. Branch thickness indicates bootstrap support. Number of unique flg22 peptide variants for each genus are listed in grey. **b,** Frequency of flg22 variants characterized in this study in comparison to all flg22 variants mined in^18^. **c,** WebLogo and sequence alignment of flg22 from plant pathogenic bacteria, colored bars represent different bacterial classes. Pae = P*seudomonas aeruginosa*, Atu = *Agrobacterium tumefaciens*, Rrh = *Rhizobium rhizogenes*, Rso = *Ralstonia solanacearum*, Ddi = *Dickeya dianthicola*, Eam = *Erwinia amylovora*, Pvi = *Pseudomonas viridiflava*, Xfr = *Xanthomonas fragariae*, Xor = *Xanthomonas oryzae*. **d,** Normalized ROS production after flg22 treatment (100nM). NbFLS2 perception was analyzed in wild-type *N. benthamiana.* Other FLS2 homologs were transiently expressed in the *N. benthamiana fls2-1/2* CRISPR/Cas9 mutant. Leaf discs for ROS assays were harvested 24 hours post infiltration. Normalized ROS levels were calculated by adjusting maximum relative light unit (RLU) averages to a 0 to 100 scale, referencing controls (water and Pae flg22). Data points below a value of 10 are depicted as white bubbles. Nb = *Nicotiana benthamiana*, Tj = *Trachelospermum jasminoides*, Qv = *Quercus variabilis*. **e,** Mitogen-activated protein kinase (MAPK) phosphorylation triggered by flg22 variants (100nM). FLS2 expression occurred as described in (d) and tissue was harvested 24h post infiltration. Coomassie brilliant blue (CBB) staining indicates protein loading. All experiments have been repeated at least three times independently with similar results.

Next, the ability of FLS2 homologs from four species to perceive the 11 flg22 variants was characterized. Wild-type *Nicotiana benthamiana* was used to investigate perception of NbFLS2-1/2. The *N. benthamiana fls2-1/2* CRISPR/Cas9 mutant was used to investigate the perception of FLS2^XL^, TjFLS2, and QvFLS2 using transient expression. All FLS2 receptors could be detected by western blotting (Extended Data Fig. 2c). Using ROS as a proxy for recognition, we assayed perception profiles. FLS2 from wild-type *N. benthamiana* only recognized Pae flg22 (Fig. 1d, Extended Data Fig. 2a-b). In contrast, FLS2^XL^, TjFLS2, and QvFLS2 were able to recognize Pae flg22 in addition to other polymorphic variants (Fig. 1d, Extended Data Fig. 2a-b). FLS2^XL^ detects flg22 variants from three pathogens: *Agrobacterium tumefaciens* (*Atu*)*, Rhizobium rhizogenes* (Rrh) and *Erwinia amylovora* (Eam). TjFLS2 recognizes flg22 variants from Pae, Atu, Rrh, *Dickeya dianthicola* (Ddi), and two variants from *Ralstonia solanacearum* (Rso). QvFLS2 recognizes the variants from Rso, Ddi and *Pseudomonas viridiflava* (Pvi). No FLS2 homologs recognize flg22 variants *Xanthomonas* species. Together, these results suggest that FLS2^XL^, TjFLS2, and QvFLS2 possess distinct expanded flg22 recognition profiles.

Perception of different FLS2 homologs was assessed using an additional immune output: MAPK phosphorylation (upon stimulation with 100 nM peptide). Generally, the outcomes from the MAPK assay aligned with those from the ROS assay (Fig. 1e). However, QvFLS2 induced a moderate ROS burst but no detectable MAPK phosphorylation in response to Atu and Pvi, even with a 10x higher peptide concentration (1 µM peptide) (Fig. 1d-e, Extended Data Fig. 3). This phenomenon has previously been described as “deviant peptides”, with an inconsistent immune output between ROS assay and seedling growth inhibition^14^.

### Structure guided engineering of expanded flg22 specificity from QvFLS2

We sought to engineer FLS2 with expanded flg22 recognition by introducing expanded specificity into homologs with narrow perception. The crystal structures of the LRR-RKs FLS2 and PEPR1 (PEP1 RECEPTOR 1), have highlighted the importance of concave surface residues on the inner surface of the LRR superhelix for peptide ligand binding^8,19^. Therefore, we hypothesized polymorphic residues on the concave surface contribute to receptor perception range.

First, we employed Repeat Conservation Mapping (RCM) to identify polymorphic residues within the LRR region. QvFLS2 originates from *Q. variabilis*, a member of the Fagales order. Therefore, 42 FLS2 homologs from the Fagales order were subjected to RCM. A region with significantly lower conservation scores was identified between LRRs 12-20 on residues located on the concave surface (Fig. 2a). In contrast, LRRs 9 – 11 and 21 – 28 exhibited higher levels of conservation (Fig. 2a). Next, AlphaFold-Multimer was used to model the complex of QvFLS2 with the co-receptor NbSERK3A and flg22 variant Rso_1. We predicted the QvFLS2 residues required for interaction with Rso_1 flg22 as those within 5 angstroms (Extended Data Fig. 4b). Diverse FLS2 concave surface residues identified by RCM are also predicted to interact with both the flg22 C-terminus and co-receptor NbSERK3A.These data suggest that the expanded perception profile of QvFLS2 is determined by variations in surface residues.

**Figure 2.**
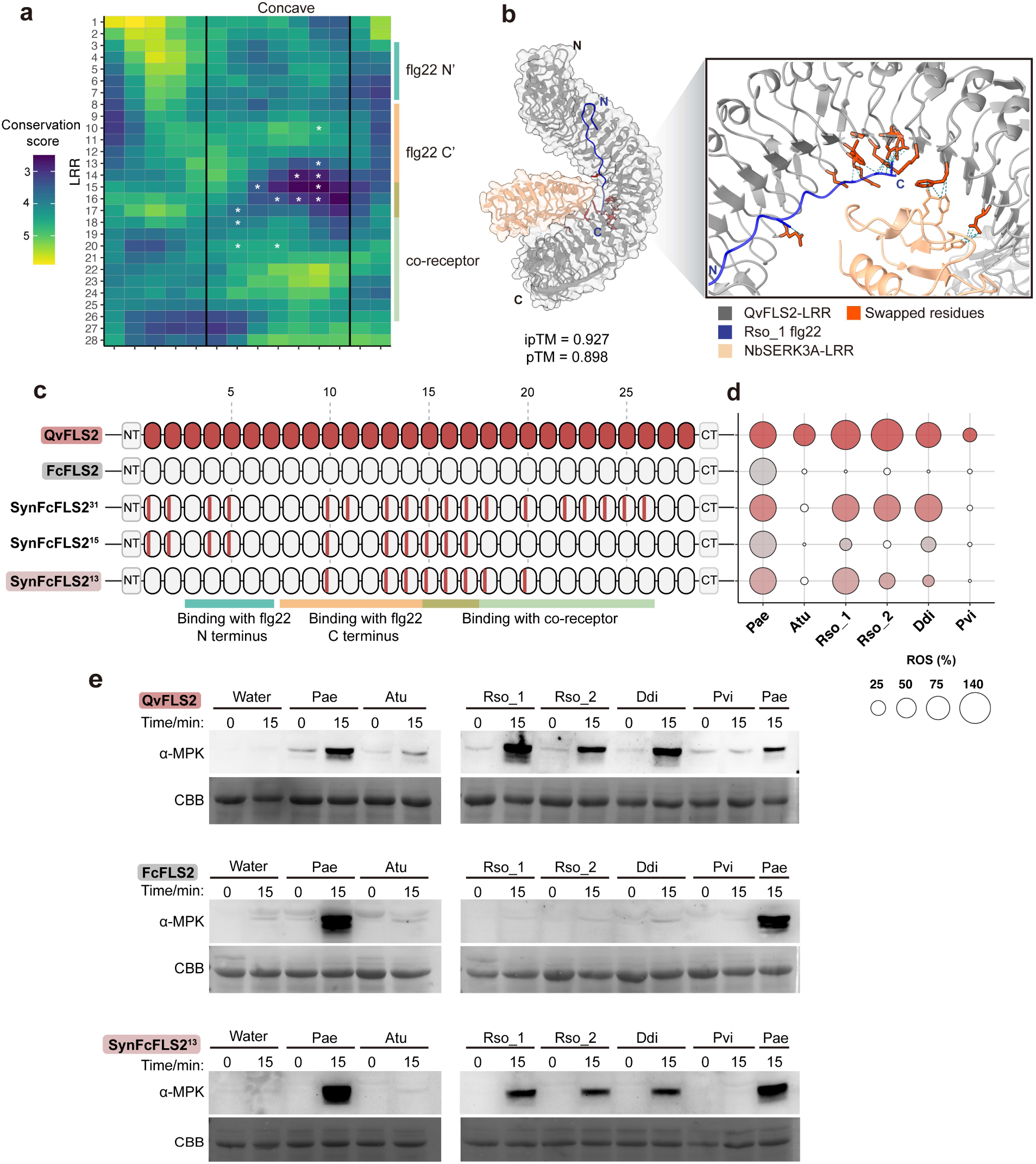
Engineering expanded ligand perception from QvFLS2 recognizing *Ralstonia* and *Dickeya* pathogens. **a,** Repeat Conservation Mapping (RCM) of 42 FLS2 homologs within the Fagales order, showing conservation levels from high (yellow) to low (dark blue). White asterisks mark the residues transferred from QvFLS2 to FcFLS2 in Synthetic FcFLS2 (SynFcFLS2^13^), which are associated with the expanded ligand detection capability. Color bars indicate the interacting regions of flg22 ligand or co-receptor. Qv = *Quercus variabilis*, Fc = *Fagus crenata*. **b,** AlphaFold model of the LRR extracellular domains of QvFLS2 and co-receptor NbSERK3A with *Ralstonia solanacearum* (Rso_1) flg22 peptide. Residues in red are associated with expanded perception as mentioned in (a). The N- and C-termini of flg22 and FLS2 are labeled. **c,** Schematic representations of Synthetic FcFLS2 (SynFcFLS2) incorporating QvFLS2 residues. Color bars within the LRR repeats indicate the occurrence of residue transfers. The number of residues swapped for each synthetic variant are superscripted after the receptor name. **d,** Normalized ROS production after treatment with flg22 variants (100nM). FLS2s were transiently expressed in the *N. benthamiana fls2-1/2* CRISPR/Cas9 mutant. Leaf discs for ROS assays were harvested 24 hours post infiltration. Normalized ROS levels were calculated by adjusting maximum relative light unit (RLU) averages to a 0 to 100 scale, referencing controls (water and Pae flg22) for each FLS2. Data points below a value of 10 are depicted as white bubbles. Pae = *Pseudomonas aeruginosa*, Atu = *Agrobacterium tumefaciens*, Rrh = *Rhizobium rhizogenes*, Rso = *Ralstonia solanacearum*, Ddi = *Dickeya dadantii*, Pvi = *Pseudomonas viridiflava*. **e,** Phosphorylation of MAPK induced by flg22 variants (100 nM). FLS2 expression occurred as described in (d) and tissued was harvested 24h post infiltration. CBB = protein loading. All experiments have been repeated independently at least three times with similar results.

To engineer expanded perception, we identified a close homolog of QvFLS2 within the Fagaceae family, *Fagus crenata* (FcFLS2). FcFLS2 shares 87.48% amino acid sequence identity with QvFLS2 but only recognizes Pae flg22 (Fig. 2c-d). Next, we aimed to determine if complete transfer of concave surface residues is sufficient to confer expanded ligand specificity. With 31 residue swaps, SynFcFLS2^31^ exhibited a similar ROS response magnitude against Rso_1, Rso_2, and Ddi flg22 as QvFLS2 (Fig. 2c-e, Extended Data Fig. 4a). However, it failed to recognize Atu and Pvi flg22, likely due to QvFLS2’s incomplete response to these variants (Fig. 2c-e, Extended Data Fig. 3).

We then designed SynFcFLS2^15^ that incorporates 15 residue swaps for QvFLS2 concave surface residues that are predicted to primarily bind the entire Rso_1 flg22 region. All designs focused on residues that differ in hydrophobicity, charge, and polarity. SynFcFLS2^15^ encompassed residue swaps from LRRs 1-17 (Fig. 2c, Extended Data Fig. 4b). SynFcFLS2^15^ exhibited a reduction in ROS response magnitude for Rso_1, Rso_2, and Ddi flg22, indicating diverse co-receptor binding residues are important for expanded recognition (Figure 2c). Next, we generated SynFcFLS2^13^, incorporating QvFLS2 concave surface residues mainly in the flg22 C-terminus and co-receptor binding regions (Fig. 2b). Using ROS as an output, SynFcFLS2^13^ maintains a comparable response magnitude against Rso_1 flg22, but reduced response to Rso_2 flg22 and Ddi flg22 (Fig. 2c-e, Extended Data Fig. 4a). Rso_2 flg22 and Ddi flg22 were still able to induce MAPK in plants expressing SynFcFLS2^13^ (Fig. 2e). These data demonstrate rational receptor engineering of expanded ligand specificity by focusing on comparative analyses, protein modeling and amino acid properties.

### Structure guided engineering of expanded flg22 specificity from FLS2^XL^

Next, we sought to engineer the expanded flg22 specificity from FLS2^XL^ to its tandem repeat paralog VrFLS2, which shares 82.07% protein sequence identity, using a similar approach. VrFLS2 can recognize Pae, Ddi, and Eam flg22 (Fig. 3d, Extended Data Fig. 5a). Due to the limited diversity of sequenced genomes in the Vitales order, the diversity and conservation patterns on the RCM plot are less distinct than for QvFLS2 (Fig. 2a, 3a). Nevertheless, we observed a similar region with lower conservation scores within LRRs 10 – 20, and highly conserved regions at LRRs 7 – 10 and 21 – 24. Complete LRR repeat swaps have demonstrated that FLS2^XL^ LRRs 12-18 are important for mediating Atu flg22 perception, aligning well with our RCM result^15^. Like QvFLS2, diverse FLS2^XL^ concave surface residues map to the flg22 C-terminus and co-receptor binding regions based on AlphaFold-Multimer modeling (Extended Data Fig. 5).

**Figure 3.**
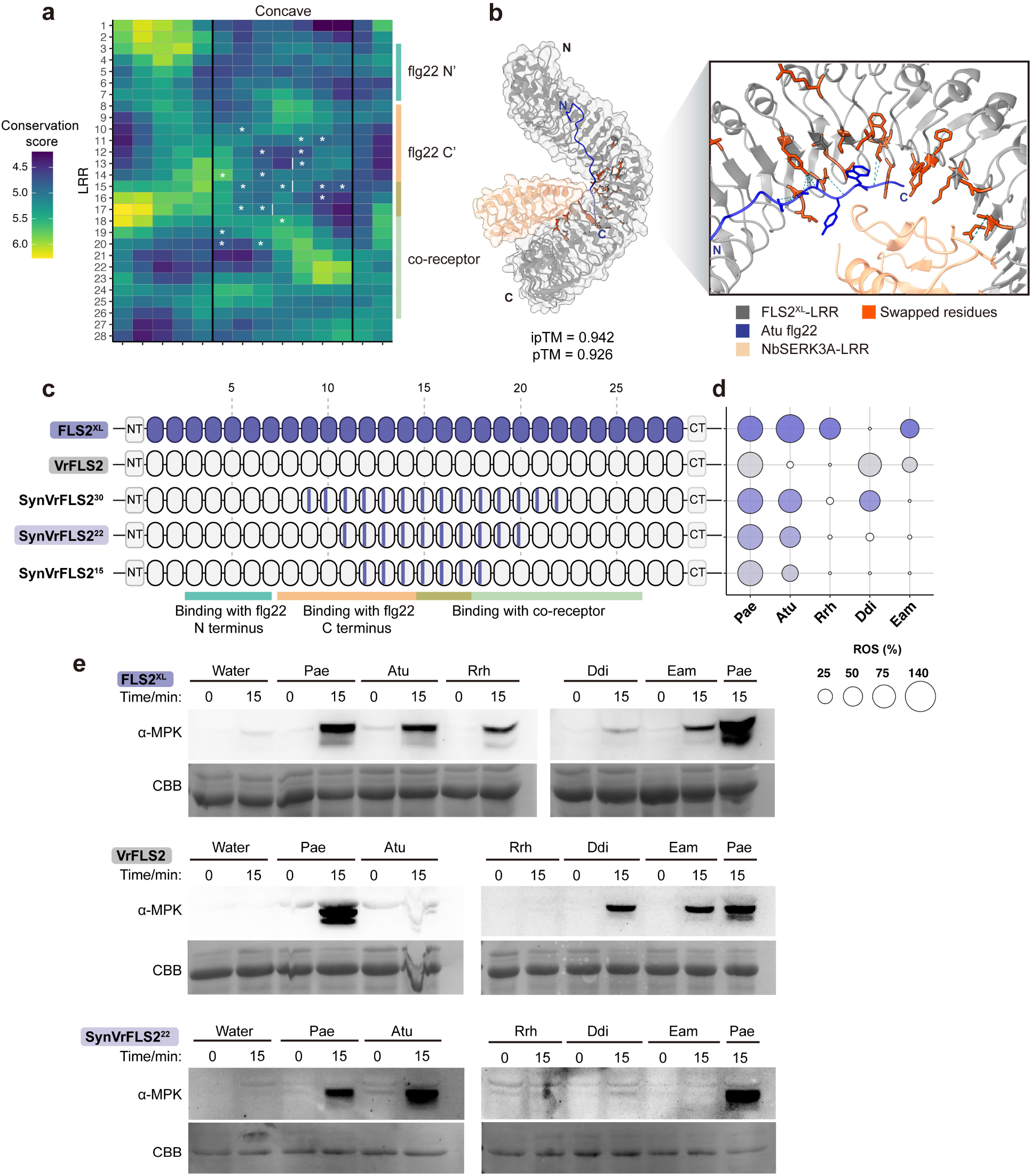
Engineering expanded ligand perception from FLS2^XL^ recognizing *Agrobacterium*. **a,** Repeat Conservation Mapping (RCM) of 14 FLS2 homologs within the Vitales order, showing conservation levels from high (yellow) to low (dark blue). White asterisks and bars mark the residues transferred from FLS2^XL^ to VrFLS2 in Synthetic VrFLS2 (SynFcFLS2^22^), which are associated with the expanded ligand detection capability. Color bars indicate the interacting regions of flg22 ligand or co-receptor. Vr = *Vitis riparia*. **b,** AlphaFold model of the LRR extracellular domains of FLS2^XL^ and co-receptor NbSERK3A with *Agrobacterium tumefaciens* (Atu) flg22 peptide. Residues in red are associated with expanded perception as mentioned in (a). The N- and C-termini of flg22 and FLS2 are labeled. **c,** Schematic representations of Synthetic VrFLS2 (SynVrFLS2) incorporating FLS2^XL^ residues. Color bars within the LRR repeats indicate the occurrence of residue transfers. The number of residues swapped for each synthetic variant are superscripted after the receptor name. **d,** Normalized ROS production after treatment with flg22 variants (100nM). FLS2s were transiently expressed in the *N. benthamiana fls2-1/2* CRISPR/Cas9 mutant. Leaf discs for ROS assays were harvested 24 hours post infiltration. Normalized ROS levels were calculated by adjusting maximum relative light unit (RLU) averages to a 0 to 100 scale, referencing controls (water and Pae flg22) for each FLS2. Data points below a value of 10 are depicted as white bubbles. Pae = *Pseudomonas aeruginosa*, Atu = *Agrobacterium tumefaciens*, Rrh = *Rhizobium rhizogenes*, Ddi = *Dickeya dianthicola*, Eam = *Erwinia amylovora*, Pvi = *Pseudomonas viridiflava*. **e,** Phosphorylation of MAPK induced by flg22 variants (100 nM). FLS2 expression occurred as described in (d) and tissued was harvested 24h post infiltration. CBB = protein loading. All experiments have been repeated independently at least three times with similar results.

To explore whether expanded recognition could be engineered, we generated three synthetic variants in the VrFLS2 backbone. Each SynVrFLS2 variant comprised residue swaps on the concave surface within regions predicted to interact with both the flg22 C-terminus and co-receptor NbSERK3A (Fig. 3c, Extended Data Fig. 5b). Consistent with observations from QvFLS2 engineering, a gradual reduction in the number of residue swaps led to decreased response magnitude of Atu flg22 (Fig 3c-e, Extended Data Fig. 5a). Notably, by swapping 22 residues, SynVrFLS2^22^ effectively recognizes Atu flg22, triggering both a ROS burst and MAPK phosphorylation (Fig 3b-e). Collectively, these findings demonstrate that two FLS2 receptors, from diverse plant orders, can be used to engineer expanded ligand perception using similar approaches.

### Variations in surface residue properties correlate with ligand specificity

Diverse amino acid properties influence the binding potential of protein-peptide interactions^20,21^. We demonstrated that residues within the concave surface are critical for mediating perception of diverse flg22 peptides (Fig. 2-3). Next, we investigated how amino acid properties contribute to binding potential and perception range. Using an unbiased approach, global values of exposed residues on the concave surface from eight FLS2 homologs with canonical or expanded ligand perception were calculated. This analysis spanned 44 different amino acid properties and scales.

First, a principal component analysis was performed to identify which amino acid properties correlate with receptor perception profiles (Extended Data Fig. 6a). Dimension one explained the greatest proportion of the variance (97.1%) across all FLS2 homologs, but the relative contribution per homologs was similar (Fig. 4a). Assessing the relative contribution of individual chemical parameters for dimension 1 revealed bulkiness and hydrophobicity (using the Manavalen scale) predominately contributed (Fig. 4b). Dimension two accounted for 2.8% of the total variance, with charge being the primary contributing factor (Fig. 4b). Strikingly, the relative contribution of dimension two varied greatly across FLS2 homologs, with it being the strongest contributor to variance for FLS2^XL^ and QvFLS2 (Fig. 4a).

**Figure. 4.**
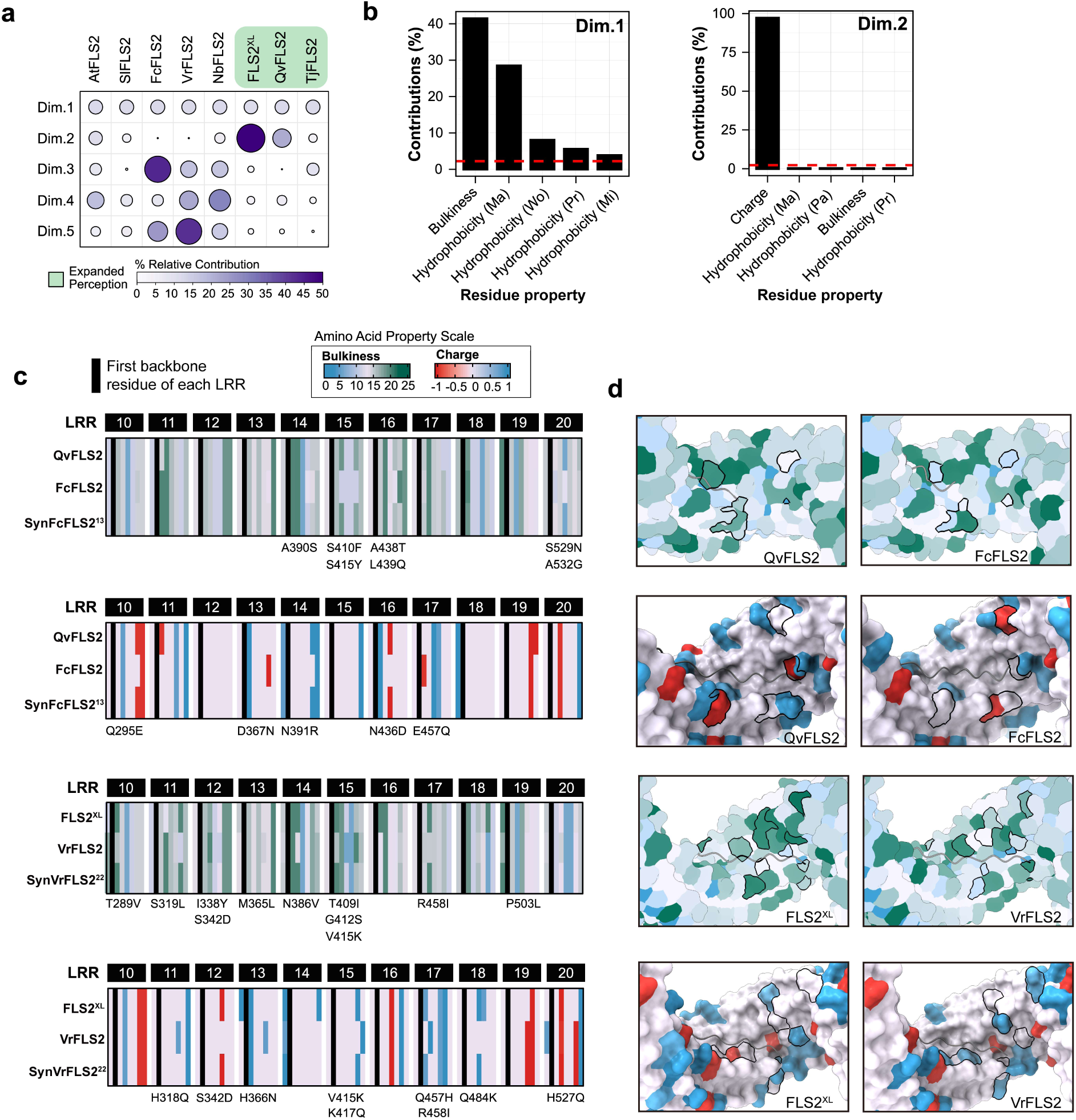
Principal component analyses identifies amino acid properties contributing to expanded ligand perception. **a,** Principal component analysis (PCA) of FLS2 homologs and their associated residue properties. First five dimensions are displayed (Dim.1 through Dim.5) and the relative contributions across each FLS2 homolog**. b,** Highest ranked contribution of chemical residue variables for top two PCA dimensions**. c,** Heatmap of bulkiness (top) and amino acid charge (bottom) along exposed residues of the LRR for QvFLS2, FcFLS2, SynFcFLS2^13^ (Fig. 2); FLS2^XL^, VrFLS2, SynVrFLS2^22^ (Fig. 3). Residues swaps within synthetic variants which display differential bulkiness and/or charge are denoted (Δ bulkiness > ±2, Δ charge > ± 0.5). Surface exposed residues are defined as “xLxxLxLxxNx” of each LRR. **d,** AlphaFold model of the LRR regions of FLS2^XL^, VrFLS2, QvFLS2 and FcFLS2 with residues colored by bulkiness or charge as labeled in c. Swaps of interest are outlined in black.

Next, we investigated differences in bulkiness and charge values of concave surface-exposed residues between FLS2 homologs with canonical, expanded, and engineered expanded perception (Fig. 4c-d, Extended Data Fig. 6b-c). We focused on LRRs 10-20, which bind flg22’s C-terminus, the SERK co-receptor, and are sufficient for engineering expanded perception. Bulky and charged residues differ in location for QvFLS2 and its canonical counterpart, FcFLS2 (Fig. 4a-b). The residue profile of SynFcFLS2^13^ closely matched QvFLS2 for bulkiness within LRRs 14-20 and charge within LRRs 10-17 (Fig. 4c, Supplementary Table 1). Overall, FLS2^XL^ carried more bulky residues than its canonical counterpart VrFLS2 between LRRs 10-19 (Fig. 4c). The bulky residue profile of SynVrFLS2^22^ closely matched that of FLS2^XL^ (Fig. 4c, Supplementary Table 1). Alterations in charge between FLS2^XL^ and VrFLS2 generally occurred within LRRs 11-18. The charge profile of SynVrFLS2^22^ closely matched that of FLS2^XL^ (Fig. 4c). Broadly, pairing amino acid properties with protein modeling will enable more precise modifications to adjust epitope recognition profiles.

### Evaluating AlphaFold3’s accuracy in predicting flg22 recognition

Employing confidence scores from AlphaFold has facilitated the identification of protein binding interfaces^22^. Although most research has focused on structured proteins, the efficacy of AlphaFold in recognizing peptide ligands remains less explored. We evaluated the accuracy of AlphaFold3 for predicting FLS2-flg22 binding as a proxy for recognition coupled with experimental data from this study and others. We modeled 221 FLS2-flg22 complexes, encompassing 40 FLS2 homologs and 2-97 natural flg22 variants per receptor (Supplementary Data 1). The average ipTM (interface predicted template modelling) and pTM (predicted template modelling) values were extracted from three independent modeling attempts as an index for predicting flg22 recognition.

Overall, AlphaFold3 can differentiate perception of FLS2-flg22 through average ipTM scores, with non-perceiving pairs exhibiting significantly lower ipTM values (Fig. 5a). Receiver Operating Characteristic (ROC) analysis is commonly used to assess model performance, with an area under the curve value of one indicating complete accuracy^23^. ROC analyses indicates that AlphaFold3 has a near perfect accuracy for predicting flg22 perception outcomes with AtFLS2, with an area under the curve of 0.99 (Fig. 5b). AlphaFold3 exhibits moderate accuracy for predicting perception of other FLS2 homologs (area under the curve = 0.77) (Fig. 5b). To maximize overall accuracy, we established an optimal ipTM threshold of 0.83 (Fig. 5c). When the ipTM threshold was set to 0.83 for all data, AlphaFold3 achieves an overall accuracy rate of 84.61%, successfully predicting 94.04% of overall non-perceiving pairs and 64.29% of perceiving pairs (Fig. 5c-d). AlphaFold3 consistently positions the peptide ligand correctly within the binding pocket, even with impossible ligands such as csp22 (Fig. 5e). To investigate if accuracy was impacted by flg22 variants, we analyzed accuracy using the FLS2 homologs and 10 flg22 variants investigated in this study (Fig. 5f). Notably, flg22 variants’ similarity to Pae flg22, included in the AtFLS2/flg22/BAK1 crystal structure, did not significantly impact prediction accuracy (Fig. 5f). In summary, AlphaFold3 demonstrates promise in predicting the recognition of peptide ligands by FLS2. Coupling AlphaFold modeling with amino acid properties could accelerate the accurate identification of PRRs with novel recognition specificity.

**Figure 5.**
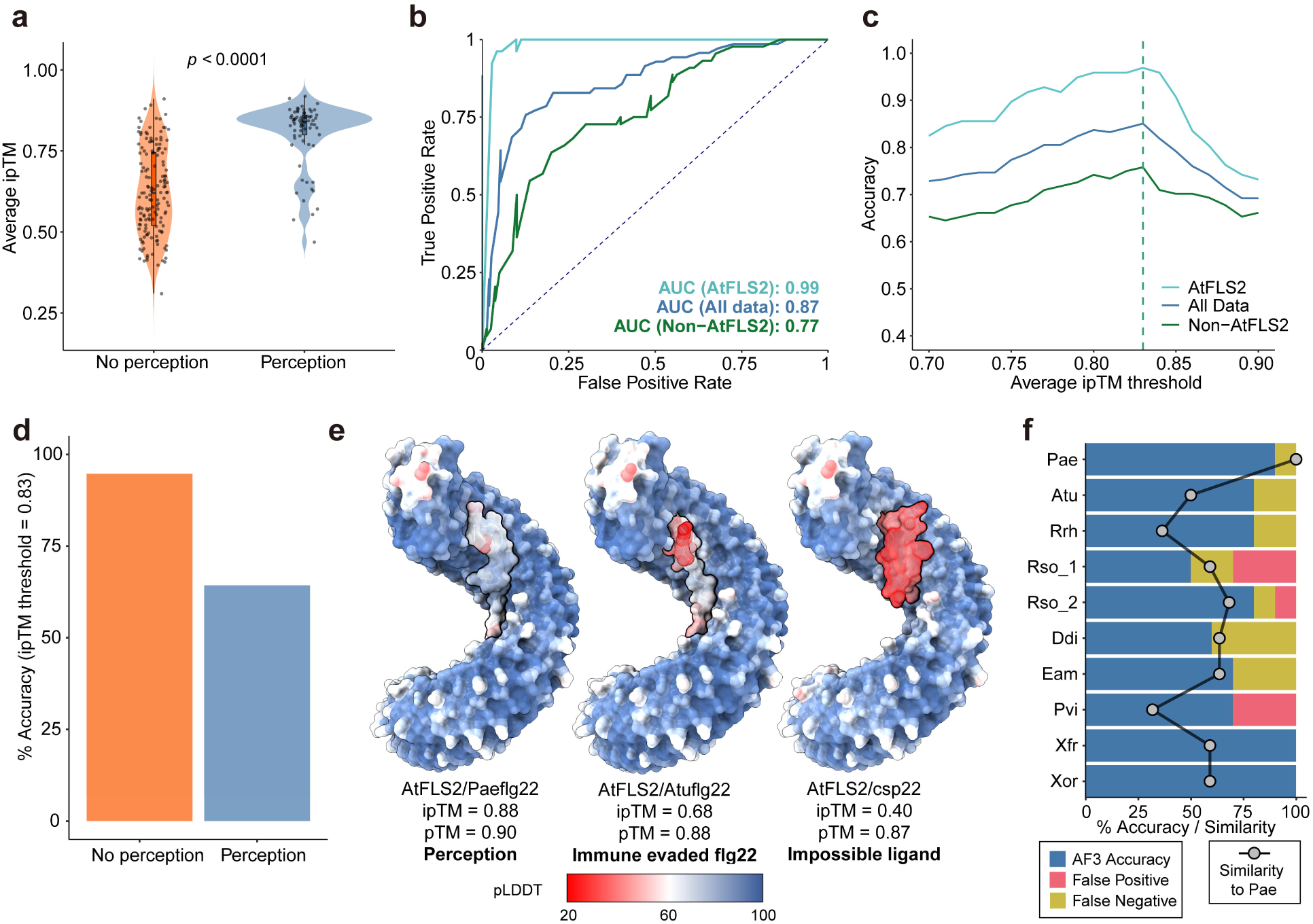
Identifying AlphaFold3 metrics for accurate prediction of flg22 recognition. **a,** Average ipTM score for each FLS2-flg22 pair whose perception has been experimentally validated (n = 221 combinations). Each dot represents the average ipTM of three independent predictions for each FLS2-flg22 pair, modelled by AlphaFold3. Wilcoxon rank-sum test was used to determine significance. The box in the boxplot represents the interquartile range (IQR), with the lower and upper hinges corresponding to the first and third quartiles, respectively. The whiskers extend to the smallest and largest values within 1.5 times the IQR from the hinges, while the central line indicates the median. **b,** Receiver operating characteristic analysis of AlphaFold3. Three groups were separately analyzed: all FLS2-flg22 pairs (blue, n = 221), only AtFLS2 (cyan, n = 97), and non-AtFLS2 (green, n = 124). The diagonal dashed line represents a random classifier, and area under the curve (AUC) values are annotated for each group, indicating the model’s performance. **c,** Accuracy curves of AlphaFold3 with different ipTM thresholds. Dashed vertical lines indicate the optimal threshold points for each group where accuracy is maximized with the ipTM threshold = 0.83. All FLS2-flg22 pairs: accuracy = 85.07%; only AtFLS2 pairs: accuracy = 96.91%; non-AtFLS2: accuracy = 75.81%. **d,** The accuracy of AlphaFold3 using a 0.83 ipTM threshold for all pairs that can perceive or not perceive flg22 ligands. **e,** AlphaFold3 models of AtFLS2 (*Arabidopsis thaliana*) with flg22 from Pae that is perceived, the flg22 variant from Atu that is not perceived, and the impossible ligand csp22 from cold shock protein. High pLDDT (predicted local distance difference test) indicates a higher confidence and more accurate prediction. **f,** Accuracy of AlphaFold3 for predicting immunogenicity of flg22 variants selected in this study (n = 10 receptors x 10 flg22 variants). Dots indicate the percentage similarity of each flg22 variant to the canonical Pae flg22.

### Evolutionary Analysis Identifies Diversifying Selection on FLS2 Concave Surface Residues

Plant immune receptors are subject to diversifying selection, driving functional divergence^24–27^. Broadened perception likely provides a selective advantage in pathogen recognition and relevant residues may be under positive selection. Therefore, we investigated whether residues in FLS2’s ectodomain are under positive selection. Three distinct phylogenetic groups comprising two plant orders and one family were selected: Brassicaceae, Fagales, and Rosales. These groups were selected based on genomic resources and the presence of multiple genomes carrying a single copy of *FLS2* that is syntenic (Extended Data Fig. 7). Positive selection analysis was conducted using CodeML. Similar patterns of selection pressure were found across each group when scanning all 28 LRRs (Fig. 6a). Specifically, LRR 11 and LRR 23 were conserved (Fig. 6a). In contrast, residues with a higher dN/dS (the ratio of non-synonymous to synonymous substitution rates) value were predominantly located in LRRs 5–8 and LRRs 12–19 (Fig. 6a and b).

**Figure 6.**
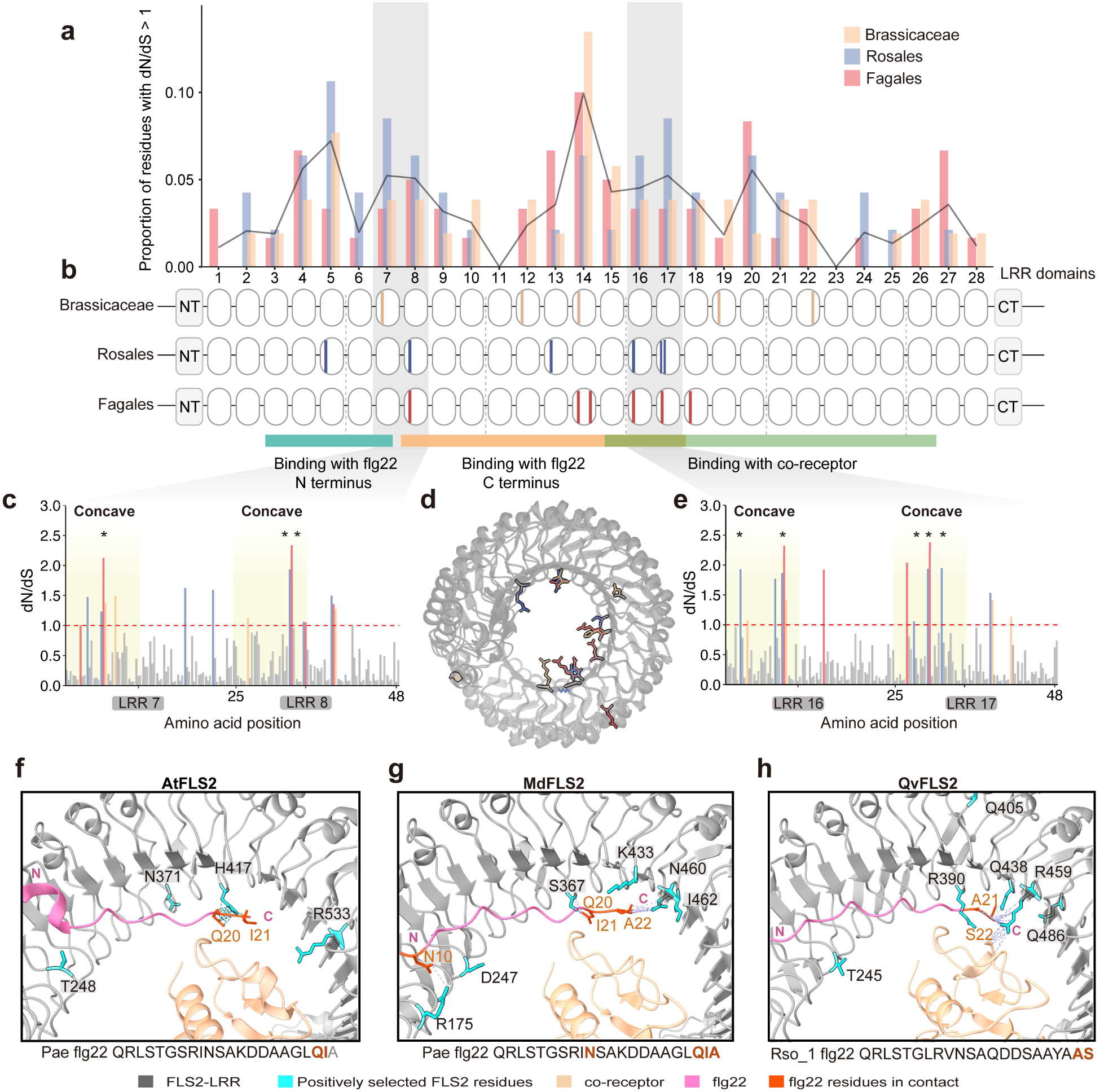
Evolutionary analysis of FLS2 from the Brassicaceae, Rosales and Fagales. **a,** Normalized proportion of FLS2 residues with dN/dS >1 across 28 LRR domains. The black line represents the average proportions from the three phylogenetic groups across each LRR domain. The dN/dS per codon is calculated using Model 8 from CodeML. **b,** Schematic representations of FLS2 from three plant groups, with color bars highlighting residues highly likely to be under positive selection with a Bayes Empirical Bayes (BEB) probability greater than 0.95. **c,** dN/dS per codon for LRR 7 and 8. Residues with dN/dS >1 are highlighted with color scheme representing Brassicaceae (peach), Rosales (slate) and Fagales (salmon). Concave surface regions are shaded with light yellow. Residues under positive selection are marked with an asterisk (BEB probability > 0.95). **d,** Overlay of the crystal structure of AtFLS2 (PDB: 4MN8) with AlphaFold models of MdFLS2 (*Malus domestica*) and QvFLS2, representing the three plant phylogenetic groups. Residues under positive selection (BEB probability > 0.95) are highlighted with the same color scheme described in b. **e,** dN/dS per codon for LRR 16 and 17. Residues with dN/dS >1 are highlighted with the same color scheme described in b. Regions on the concave surface are shaded in light yellow. Residues under positive selection are marked with an asterisk (BEB probability > 0.95). **f to h,** Structural visualization of the AtFLS2/BAK1/Paeflg22 (PDB: 4MN8) complex alongside AlphaFold models of MdFLS2/MdSERK1/Paeflg22 and QvFLS2/NbSERK3A/Rso_1flg22. FLS2 residues highly likely to be under positive selection are highlighted in cyan, while residues of the flg22 and co-receptors interacting with positively selected FLS2 residues (within a 5 Å distance) are shown in orange. The N- and C-termini of flg22 are labeled.

To identify specific residues under positive selection, the Bayes Empirical Bayes (BEB) method was used to calculate selection probabilities. Residues with a BEB probability exceeding 0.95 were considered positively selected. These residues are highlighted in Fig. 6b and mapped onto the structural models of AtFLS2, MdFLS2 (*Malus domestica*), and QvFLS2, representing the Brassicaceae, Rosales, and Fagales, respectively (Fig. 6d). Of these positively selected residues, 88.2% were located on the concave surface of the LRR, mainly near the binding pockets for the flg22 polymorphic C-terminus and the co-receptor (Fig. 6f-h). This aligns with previous findings that polymorphisms in the flg22 C-terminus are key for canonical FLS2 immune evasion (Fig. 6c to e)^15,17^. The selection analyses mirror our experimental data for QvFLS2 and FLS2^XL^ engineering. Overall, our data suggests that integrating evolutionary analysis, protein modeling, and amino acid properties is a promising strategy to identify receptors with expanded perception and pinpoint specific residues for engineering broader recognition.

## Discussion

Plants contain many PRRs, but MAMP diversity impairs receptor effectiveness. Using an interdisciplinary approach, we identified key residues along FLS2’s concave surface that enabled expanded recognition and engineered them into homologs with canonical perception. Our findings identify amino acid properties and AlphaFold3 metrics for predicting PRR recognition, providing a framework for engineering receptors to improve pathogen restriction.

Previous studies have demonstrated that exchanging sets of LRRs can alter ligand specificity of immune receptors^28–30^. Swapping LRRs 19-24 from SlFLS2 (S*olanum lycopersicum*) into AtFLS2 affected binding affinity to artificial flg22 variants^31^. In FLS2^XL^, LRR subdomain swaps with its close paralog VrFLS2 highlighted the critical role of LRRs 12-18 in mediating Atu flg22 perception^15^. These data are consistent with our findings, which demonstrate the importance of specific residues in LRRs 10-20 for conferring expanded perception (Fig. 2-3). Previous efforts to create receptor variants with expanded recognition through site-directed random mutagenesis have slightly enhanced AtFLS2 sensitivity to polymorphic flg22 at micromolar concentrations^32^. We were able to pinpoint key residues crucial for ligand specificity by focusing on differences in the chemical properties of residues on the concave surface between receptors with different perception ranges (Fig. 2-3). A recent study focused on engineering the NLR receptor Sr50 to recognize a variant of the effector ligand AvrSr50, which evades immune detection^33^. Similarly, they showed that substituting residues on the LRR concave surface with others differing in charge or polarity can broaden the receptor’s recognition capabilities^33^. These findings suggest that targeting LRR concave surface residues is a promising approach for engineering novel ligand perception across diverse LRR-containing immune receptors.

AlphaFold’s ability to predict protein-protein interactions has rapidly evolved. AlphaFold prediction of the NLR MLA3 (Mildew locus a 3) in complex with the fungal effector PWL2 (Pathogenicity toward Weeping Lovegrass 2), revealed that MLA3 shares a similar binding interface with other PWL2 targets^34^. Furthermore, this interface could be transferred to another NLR, resulting in an expanded effector recognition profile^34^. AlphaFold can predict the interactions between the Arabidopsis RK MIK2 (MALE DISCOVERER 1-INTERACTING RECEPTOR-LIKE KINASE 2) and its corresponding plant peptide ligands^35^. We used AlphaFold to pinpoint residues on the binding interface between FLS2 homologs and flg22 variants to engineer novel recognition specificities (Fig. 2-3). Predictions involving AtFLS2, which has a known crystal structure, and various flg22 variants achieved a surprisingly high accuracy of 96.91% (ipTM threshold of 0.83). In contrast, pairs excluding AtFLS2-flg22 reached 75.81% accuracy (ipTM threshold of 0.83). This indicates that obtaining additional crystal structures of receptor-ligand complexes will improve AlphaFold3’s accuracy to predict recognition. Future experimental investigations could determine if the ipTM thresholds for FLS2-flg22 can be broadly applied to other LRR-RKs.

Polymorphic ligand perception can occur through the functional diversification of a single receptor or the convergent evolution of a unique receptor with a distinct perception range. Beyond FLS2, some potato genotypes have homologs of the LRR-RK PERU (Pep-13 receptor unit) with expanded specificity against synthetic Pep-13 peptide variants^24^. Across two orders and one family, we detected similar patterns of diversifying selection on specific FLS2 LRRs, particularly those mapping to concave surface residues (Fig. 6). One region undergoing diversifying selection maps to LRRs 12-19, aligning with our engineering of expanded ligand perception (Fig 6). Overall, evidence suggests that expanded recognition specificity may be more common than previously thought.

Pathogens contain high evolutionary potential and can carry diverse MAMP epitopes that evade PRR perception^13,14,18^. Our findings reveal that all three FLS2 homologs with broadened perception recognize additional flg22 variants from important plant pathogens (Fig. 1). However, none have identical recognition profiles; it is unlikely that a single receptor could be engineered to recognize all flg22 variants, each may require distinct binding interfaces (Fig. 4).

This study highlights approaches to realize the potential for rational design of immune receptors by altering the binding interface. Coupling diversity analyses, protein modeling, and biochemical parameters with experimental data can facilitate the identification and engineering of novel receptors with desired specificity for important pathogens. Narrowing specificity to a few amino acid residues is compatible with genome editing technologies that will expedite the deployment of the engineered resistance genes and reduce regulatory burdens^36–38^.

## Material and Method

### Plant materials and growth conditions

*N. benthamiana* plants were grown in a Conviron growth chamber at 26°C with a 16-h light/8-h dark photoperiod (180 μM m^−2^ s^−1^). Thirty-day-old (non-flowering) plants were used for *A. tumefaciens* mediated transient protein expression and subsequent immunity assays. The *N. benthamiana fls2-1/2* CRISPR/Cas9 mutant line was provided by Qiang Cheng and used for transient expression of FLS2 constructs and subsequent ROS and MAPK assays^16^.

### Selection criteria for flg22 variants and peptide preparation

FliC sequences were collected from α, β, and γ-proteobacterial genomes assessed in a previous study^18^. Protein sequences were aligned via MAFFT (v7.310) using the following parameters: -- reorder --thread 12 --maxiterate 1000 –auto^39^. A maximum-likelihood tree was built from the alignment using IQ-TREE2 (v2.1.2) with the following parameters: -st AA -bb 1000 -m TEST -T AUTO^40^. Newick file was visualized using the following R packages: phangorn (v2.7.1), treeio (v1.14.4), and ggtree (v3.1.2.991). Epitope variants were selected from the previous study based on their abundance in the dataset, their diverse sequence in respect to the consensus flg22 epitope from *P. aeruginosa,* and unique C-terminal residues^18^.

Pae flg22 and Atu flg22 were synthesized by Genscript (≥95% purity, Piscataway, NJ, USA) and Agrisera (≥95% purity), respectively. All other flg22 variants were synthesized by Shanghai Apeptide (≥95% purity). The peptides were solubilized in either water or DMSO, following the manufacturer’s recommendations, details about the flg22 peptides can be found in (Supplementary Table 2).

### Cloning of FLS2s

Plasmids harboring the *FLS2^XL^* and *VrFLS2* cDNA sequences were provided by Georg Felix^15^. Plasmids containing the *TjFLS2* and *QvFLS2* genomic DNA sequences were supplied by Qiang Cheng^16^. All full-length *FLS2s* were initially cloned into the pENTR/D-TOPO backbone (Invitrogen #K2400-20) and subsequently moved into the binary destination vector pGWB514 using the Gateway LR Clonase II enzyme mix (Invitrogen #11791-100). The expression of *FLS2* constructs was driven by the 35S promoter and included a C-terminal HA tag. The synthetic FLS2 variants were reconstructed by assembling inserted regions that carry the amino acid changes into pENTR:FcFLS2 and pENTR:VrFLS2 using NEBuilder® HiFi DNA Assembly (NEB #E5520). The details of all used constructs and primers are in Supplementary Tables 3-4.

### Transient expression of FLS2

*FLS2* variants were cloned into pGWB514 and transformed via electroporation into *A. tumefaciens* C58C1. The *N. benthamiana fls2-1/2* mutant was infiltrated with *A. tumefaciens* suspensions of OD_600_ = 0.6 induced with infiltration media [10 mM MgCl2, 5 mM MES-KOH (pH 5.6) and 0.2 mM Acetosyringone] for 2 h. Twenty-four hours after infiltration, leaf disks were collected using a #1 or #5 cork borer (4 mm) for ROS assays and MAPK assay.

To visualize the expression of FLS2s, additional leaf disks were collected using a #7 cork borer at 24 hpi for protein extraction. The leaf disks were homogenized in 100-μL Laemmli buffer and boiled for 5 min. Western blotting was conducted as described above and visualized with anti HA-HRP antibody (Roche, #12013819001, Anti-HA-Peroxidase, High Affinity; 1:3000).

### ROS burst assay

Un-infiltrated wild type *N. benthamiana* (containing native *NbFLS2*) and the *Nbfls2* CRISPR/Cas 9 mutant transiently expressing *FLS2* were used for ROS assays. Leaf disks from test plants or infiltrated tissues were collected using a #1 cork borer (4 mm diameter) and floated overnight in 200 μL of deionized distilled water in a Corning Costar 96-Well White Solid Plate (Fisher #07-200-589), covered with a plastic lid to minimize evaporation. The next day, the water was replaced with 100 μL of an assay solution containing flg22 variants. This solution consisted of 20 μM L-012 (a luminol derivative, Wako Chemicals USA #120-04891), 10 mg/mL horseradish peroxidase (Sigma), and the flg22 variants. All treatments used 100 nM flg22 variants, with an equivalent amount of DMSO included for the control. Luminescence was measured using a BioTek Synergy H1 microplate reader (Agilent) with readings taken at 0.5-second intervals from each leaf disk over a one-hour period.

For each plate, each treatment included 16 leaf disks from four individual plants. The average maximum relative luminescence units (RLUs) for each plant were calculated by first determining the maximum RLU from the four leaf disks. These maximum values were then normalized, with the average RLU for water set to 0 and that for Pae flg22 set to 100,000. Across all ROS plates, the overall average maximum RLUs were calculated using all plates run for each flg22-FLS2 combination.

### MAPK induction assay

For native *NbFLS2*, WT *N. benthamiana* leaves were infiltrated with *A. tumefaciens* C58C1 carrying a pGWB514 empty vector. For other *FLS2s*, *N. benthamiana fls2-1/2* mutant leaves were infiltrated with constructs as described above.

Twenty-four hours after infiltration, three leaf disks from the infiltrated areas were collected using a #5 cork borer (9 mm diameter) and placed overnight in 1 mL of deionized water in a 24-well tissue culture plate (VWR #10062-896), covered with a plastic lid to minimize evaporation. The following day, the water was replaced with 900 μL of either water or water containing 100 nM flg22 variants. Leaf disks from each well were then collected individually at 0 and 15 minutes after treatment, quickly frozen in liquid nitrogen, and ground using pestles attached to an electric grinder (Conos AC-18S electric torque screwdriver). Protein extraction and MAPK assay specific Western-blotting was conducted as previously described^10^.

### Identification and evolutionary analysis of *FLS2* homologs

*FLS2* homologs were identified from available plant genomes in Brassicaceae, Rosales and Fagales order using standard BLASTN/BLASTP search on NCBI and Phytozome. Candidate sequences were removed if they were duplicated genes, pseudogenes, possessing major sequence gaps or incomplete FLS2 domain architecture. Sequence IDs for all *FLS2* homologs are provided in (Supplementary Data 2).

Full length FLS2 protein sequences were initially aligned using MAFFT v7.490. A maximum likelihood (ML) tree was built from the alignment using PhyML 3.3.2 with 1,000 bootstrap replicates^41^, using the best fit model selected by the Bayesian Information Criterion (BIC). Output trees were visualized on iTOL^42^.

To perform positive selection analyses, DNA sequences of *FLS2* LRR regions within the Brassicaceae, Rosales, and Fagales were aligned using MAFFT v7.490. Subsequently, ML trees of full length FLS2 proteins and the aligned *FLS2* LRR region DNA sequences from each group, were used for positive selection analysis under site-specific models using the CodeML package, part of the ETE Toolkit^43,44^. After Likelihood Ratio Test (LRT) and Bayes Empirical Bayes (BEB) calculations, Posterior mean dN/dS value per codon and positively selected residues (BEB posterior probability > 0.95) are extracted from M8.

To perform synteny analyses, selected annotated genomes from Brassicaceae, Fagales and Rosales are downloaded from NCBI database. After manually curating the annotations, MCscan from the JCVI suite was used to infer the synteny blocks within each group via all-against-all LAST^45^.

### Repeat conservation mapping

LRR domains of FLS2 proteins from the Vitales and Fagales orders were analyzed. Repeat conservation scores were calculated using the standalone version of the RCM tool^46^. Concave surface is defined as the “xLxxLxLxxNx” region of each LRR.

### Analyses of Residue properties on the concave surface

Concave surface-exposed residues are defined as the variable residues within the “xLxxLxLxxNx” region of each LRR. For each FLS2 homolog analyzed in this study, a global value of concave surface-exposed residues was calculated across 44 different amino acid properties and scales using the R packages Peptides (v2.4.5) and alakazam (v1.3.0)^21^. Using the R packages FactoMineR (v2.11) and factoextra (v1.0.7), a principal component analysis was conducted on the matrix and dimension one through five was visualized within the corrplot (v0.92) package. To illustrate the impact of top correlated residue properties with possible ligand perception specificity, the bulkiness and charge (at pH 5.4) values were calculated along the surface-exposed residues for two canonical, two expanded, and two synthetic variants and plotted as a heatmap using ComplexHeatmap (v2.6.2)^47^. Finally, FLS2 homologs of interest were modeled in ChimeraX with residue colored by either bulkiness or charge values.

### Structural modeling and visualization

Protein structure complex with the LRR domain of FLS2 homologs with co-receptor and flg22 variants were predicted through ColabFold (v1.5.5) using pdb100 template mode^48^. Sequence alignment was generated via MMSeq2 with unpaired-paired mode. Models with ipTM > 0.8 were saved for further analysis. FLS2 residues that are within 5 angstroms of the flg22 ligand and the co-receptor were designated as candidate interface residues to capture potential atomic contact.

For benchmarking AlphaFold3’s flg22 perception prediction, protein sequences of flg22 variants and selected FLS2-LRRs were input into AlphaFold3 Server to model the different combinations of FLS2-flg22 complex^49^. Each combination was modeled three times independently. After validating the AlphaFold3 result with experimental data, The best ipTM threshold value to achieve the highest accuracy for AF3-based prediction is 0.83. All protein structures were visualized and analyzed in Chimera-X and PyMOL.

## Data Availability

All raw data underlying each figure is available on Zenodo (10.5281/zenodo.13738226). R scripts used for data analysis and plotting are accessible on GitHub Repository (https://github.com/jerrytli/FLS2_engineering). All plasmids generated in this study were deposited to Addgene (226412 - 226420) for public distribution upon publication.

## Acknowledgements

We would like to thank Andrew Bent (University of Wisconsin-Madison) for accessing and assistance with the RCM tool. We would like to thank Alex Zanella Zaccaron for assistance with microsynteny analysis. G.C., T.L. and E.J.B. were supported by a grant from the NIH (R35GM136402). T.L. was partially supported by the China Scholarship Council no. 201906300032. D.M.S was supported by a USDA-NIFA 2021-67034-35049.

## Author Contributions

T.L. and G.C. conceived and designed the study. T.L. performed all experiments unless indicated below, including structural modeling, most biochemical experiments and FLS2 diversity analyses. E.J.B. performed MAPK and receptor expression experiments. H.S. assisted in AlphaFold3 modeling. D.M.S. performed bioinformatic analyses on flg22 epitopes and FLS2-flg22 residue properties. D.M.P. assisted in evaluating structural models. T.L. and G.C wrote the manuscript, and all authors were involved in editing.

---

**Extended Data Figure 1:**
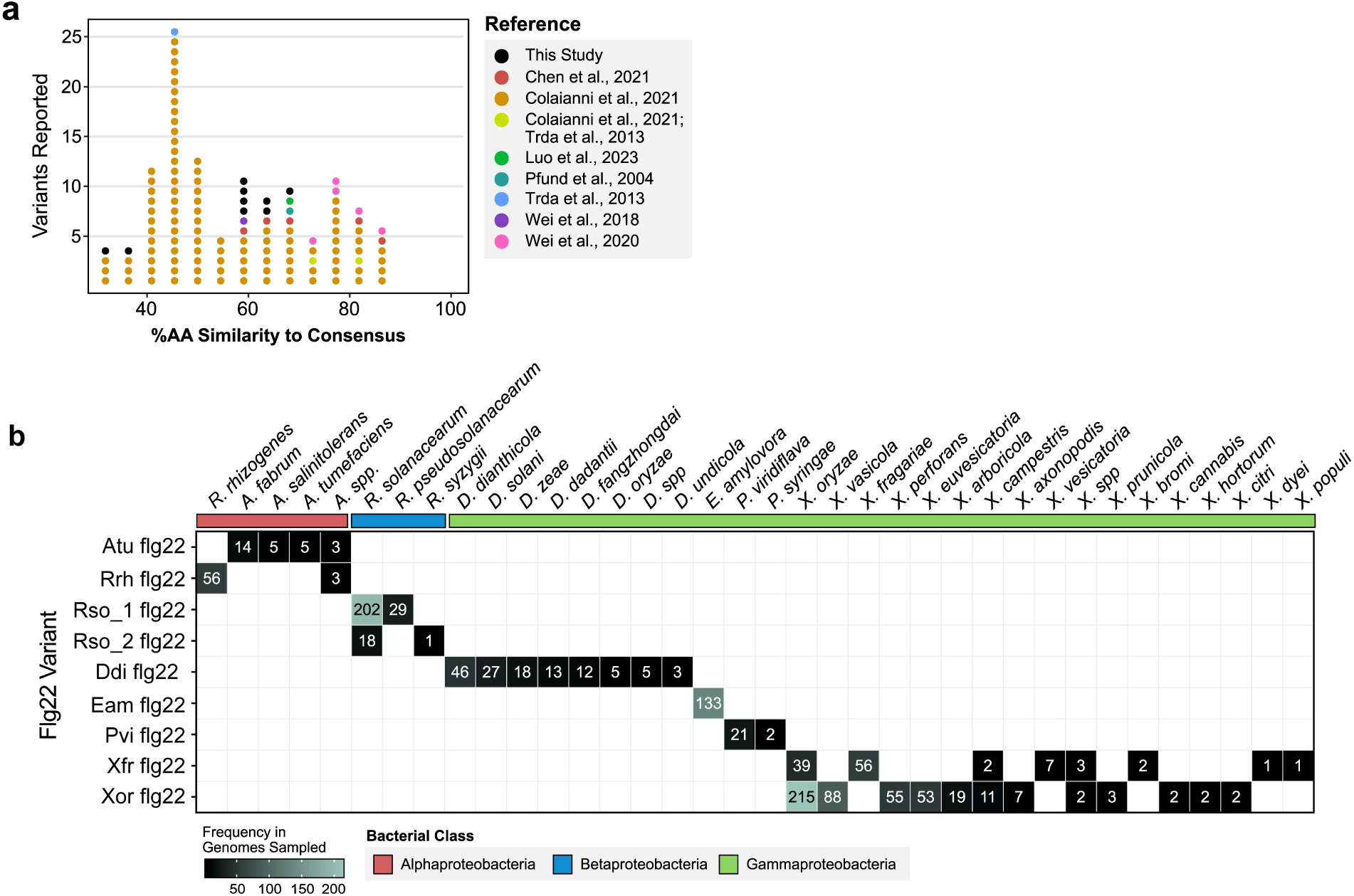
Diverse flg22 variants selected in this study. **a,** Percent amino acid (%AA) similarity of flg22 variants in comparison to the consensus selected in this study and past research. **b,** Frequency of selected flg22 variants tested in respect to their species designation from the genomes sampled.

**Extended Data Figure 2:**
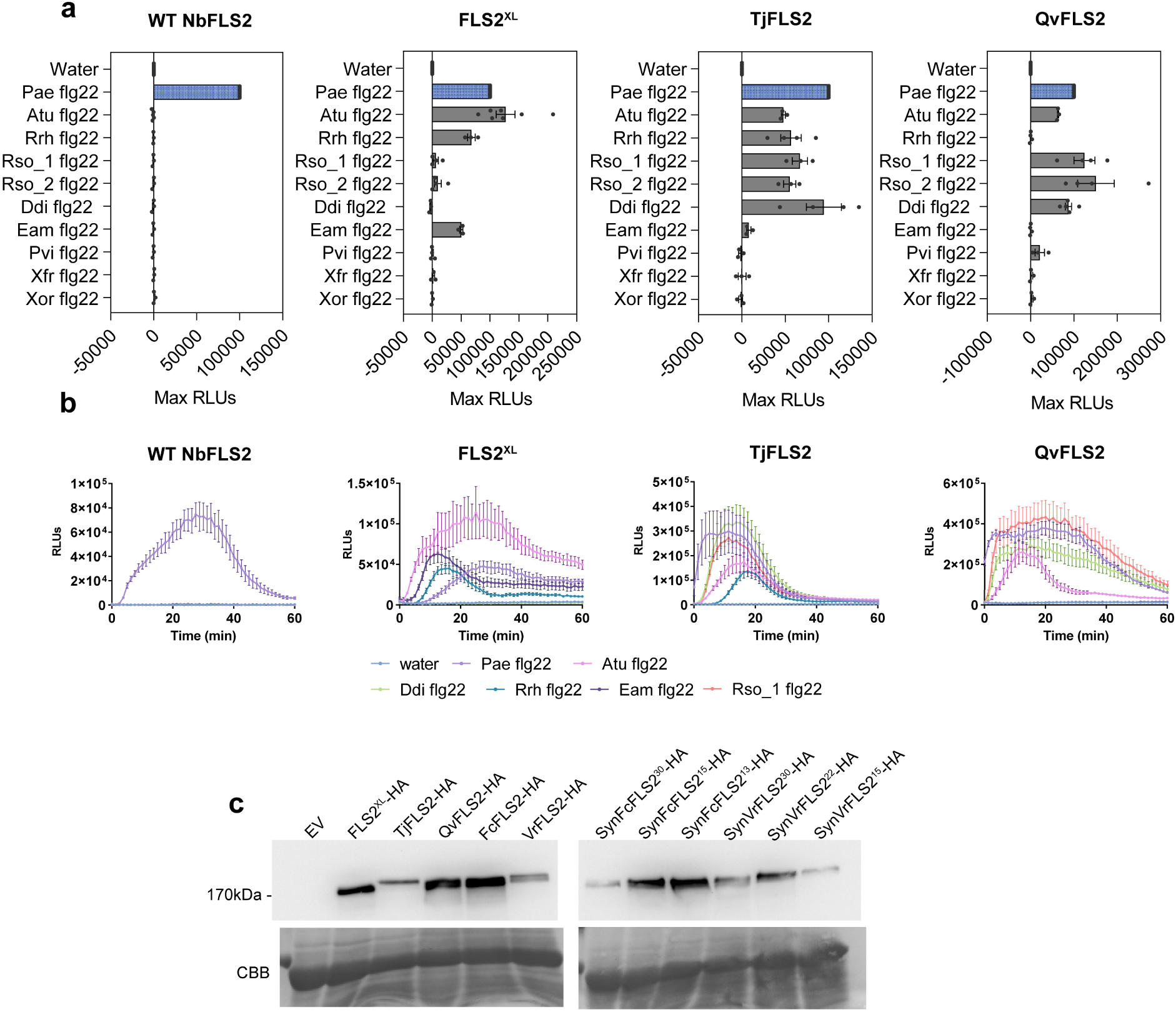
Perception of flg22 variants. **a,** Normalized ROS induction of flg22 variants (100 nM concentration) in WT *N. benthamiana* and the *N. benthamiana* fls2-1/2 expressing *FLS2^XL^*, *TjFLS2* and *QvFLS2*. Each data point represents an average max relative light unit (RLU) from 4 plants with 4 leaf disks per plant. The values of each plate were normalized by adjusting maximum relative light unit (RLU) averages to a 0 to 100000 scale, referencing negative and positive controls (water and Pae flg22). Error bars = SEM. **b,** ROS curves for a subset of flg22 variants on various FLS2 expressing plants. Error bars = SEM. All peptides were tested at 100 nM. **c,** Expression of FLS2 homologs and synthetic variants in this study.

**Extended Data Figure 3:**
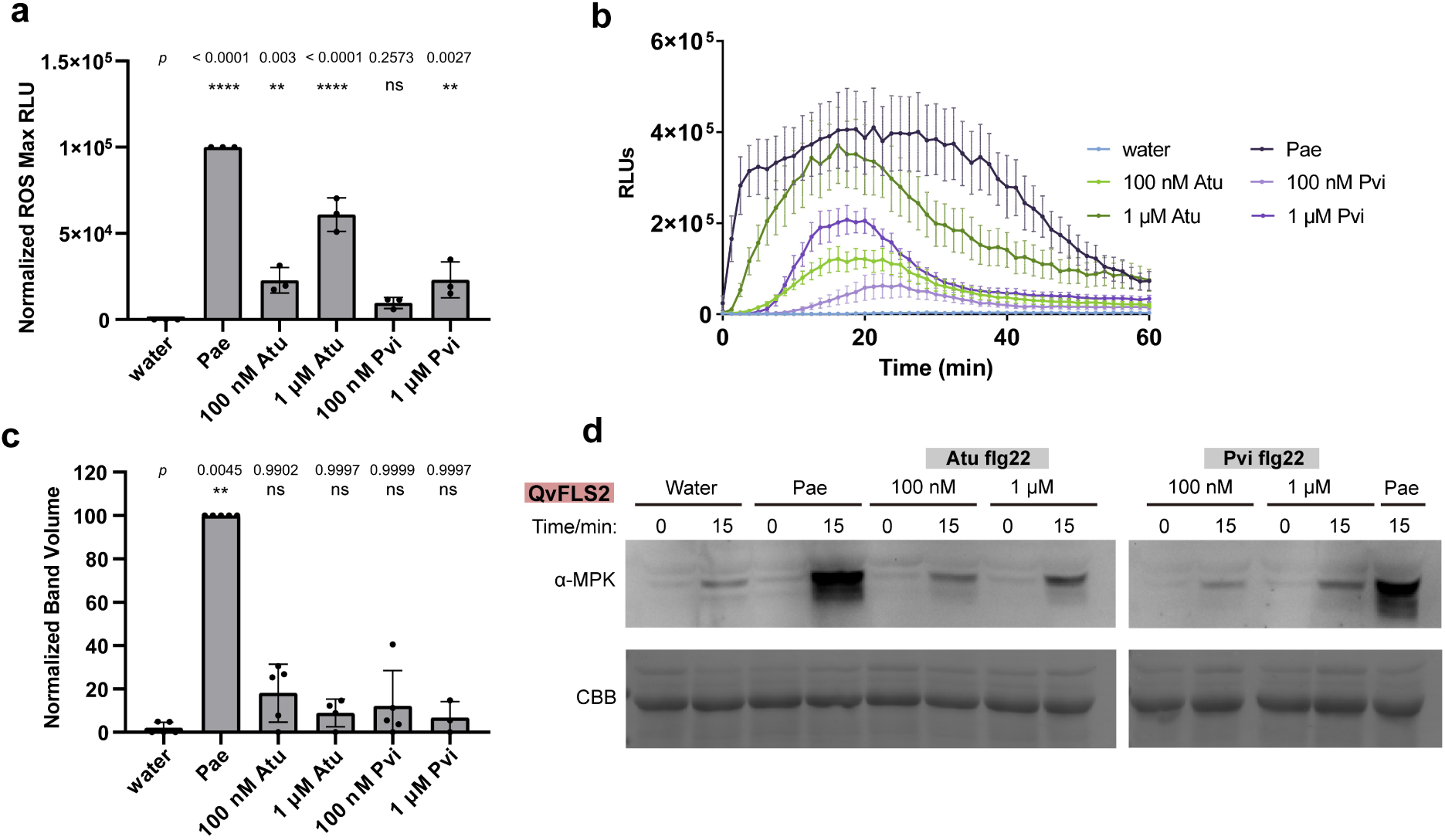
Deviant flg22s induce ROS but not MAPK. **a,** Normalized ROS induction of flg22 variants (100 nM concentration) in *N. benthamiana fls2-1/2* expressing QvFLS2. Each data point represents an average max relative light unit (RLU) from 4 plants with 4 leaf disks per plant. The values of each plate were normalized by adjusting maximum relative light unit (RLU) averages to a 0 to 100000 scale, referencing negative and positive controls (water and Pae flg22). A one-way ANOVA and Dunnett’s multiple comparison were used to determine significance using the non-scaled data, *p* values from left to right are marked on top. Error bars = SD. **b,** ROS curves for two different concentrations (100 nM and 1 μM) of Atu and Pvi. Error bars = SD. **c,** Normalized phosphorylated MAPK abundance. Bands from all treatments at 15 mins were normalized to Pae 15 mins. A one-way ANOVA and Dunnett’s multiple comparison were used to determine significance using the non-scaled data, *p* values from left to right are marked on top. Error bars = SD. **d,** Phosphorylation of MAPK induced by flg22 variants in the *N. benthamiana fls2-1/2* expressing QvFLS2. CBB = protein loading.

**Extended Data Figure 4:**
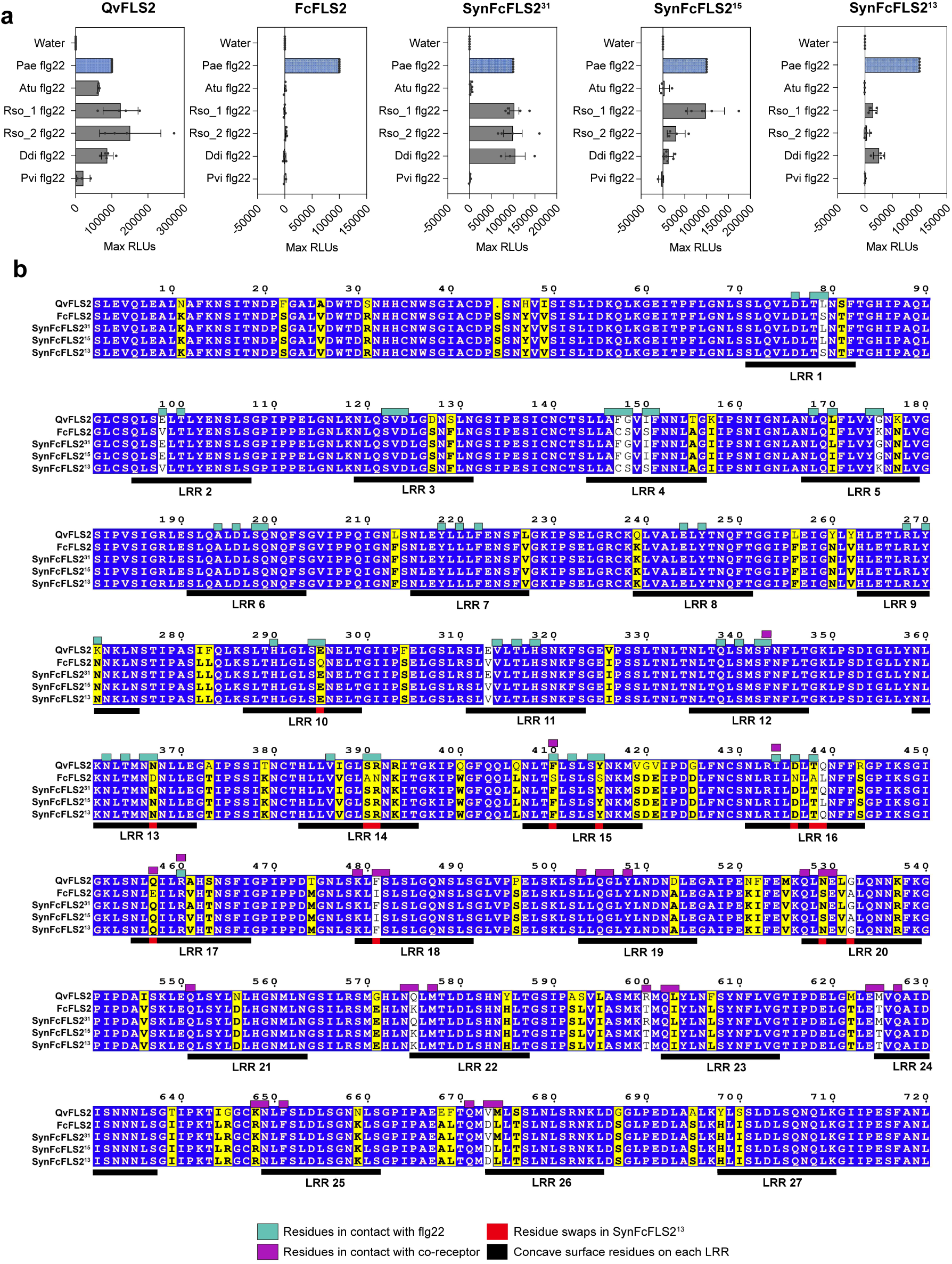
Engineering QvFLS2 specificity. **a,** Normalized ROS induction of flg22 variants (100 nM) in *N. benthamiana fls2-1/2* expressing QvFLS2, FcFLS2, and synthetic FcFLS2s. Each data point represents an average max relative light unit (RLU) from 4 plants with 4 leaf disks per plant. The values of each plate were normalized by adjusting maximum relative light unit (RLU) averages to a 0 to 100000 scale, referencing negative and positive controls (water and Pae flg22). **b,** Protein sequence alignment for LRR domains of five FLS2 variants. Concave surface regions of each LRR are highlighted with dark bars. Residues on the interacting surface are determined using the AlphaFold model of QvFLS2/NbSERK3A/Rso_1flg22 (< 5 Å). Residues in contact with flg22 are highlighted in cyan above the alignment, while residues in contact with the co-receptor are highlighted in purple. Residues swapped for SynFcFLS2^13^ are highlighted with red blocks.

**Extended Data Figure 5:**
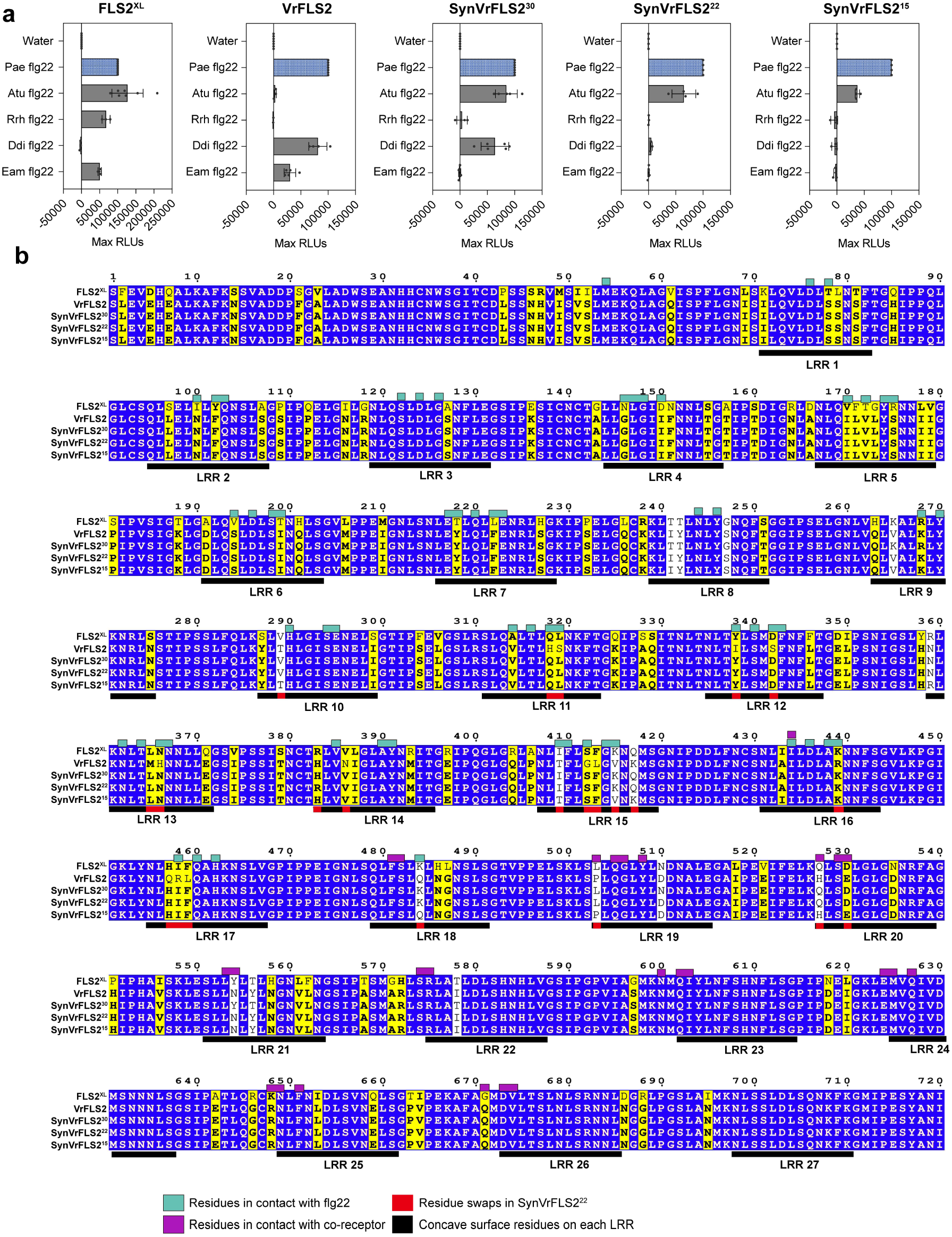
Engineering FLS2^XL^ specificity. **a,** Normalized ROS induction of flg22 variants (100 nM) in *N. benthamiana fls2-1/2* expressing FLS2^XL^, VrFLS2, and synthetic VrFLS2s. Each data point represents an average max relative light unit (RLU) from 4 plants with 4 leaf disks per plant. The values of each plate were normalized by adjusting maximum relative light unit (RLU) averages to a 0 to 100000 scale, referencing negative and positive controls (water and Pae flg22). **b,** Protein sequence alignment for LRR domains of five FLS2 variants. Concave surface regions of each LRR are highlighted with dark bars. Residues on the interacting surface are determined using the AlphaFold model of FLS2^XL^/NbSERK3A/Atuflg22 (< 5 Å). Residues in contact with flg22 are highlighted in cyan above the alignment, while residues in contact with the co-receptor are highlighted in purple. Residues swapped for SynVrFLS2^22^ are highlighted with red blocks.

**Extended Data Figure 6.**
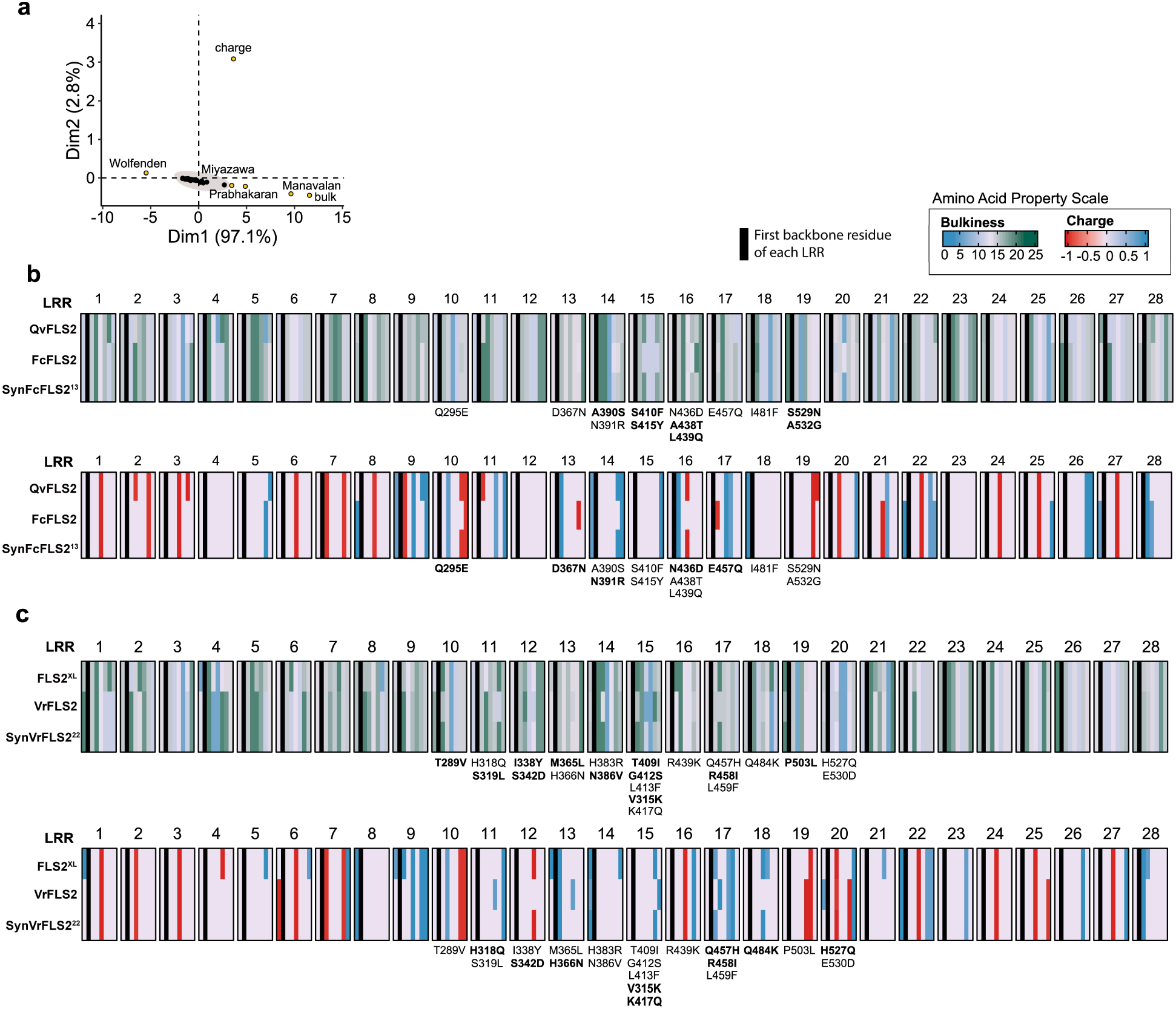
Principal component analyses identify amino acid chemical properties contributing to differential ligand perception. **a,** Principal component analysis (PCA) of FLS2 homologs and their associated residue chemistries. Scatterplot of contribution of chemical residue variables for top two PCA dimensions. Amino acid properties labeled in yellow display the highest variance and those in black display little variance between FLS2 homologs. **b,** Heatmap of bulkiness (top) and amino acid charge (bottom) along exposed residues of the LRR across QvFLS2, FcFLS2, and SynFcFLS2^13^. All swapped residues in SynFcFLS2^13^ are labeled below the heatmaps. Swapped Residues with substantial change of certain chemical parameter are highlighted in bold (Δ bulkiness > ±2, Δ charge > ± 0.5). **c,** Heatmap of bulkiness (top) and amino acid charge (bottom) along exposed residues of the LRR across FLS2^XL^, VrFLS2, and SynVrFLS2^22^. All swapped residues in SynVrFLS2^22^ are labeled below the heatmaps. Swapped Residues with substantial change of certain chemical parameter are highlighted in bold (Δ bulkiness > ±2, Δ charge > ± 0.5).

**Extended Data Figure 7:**
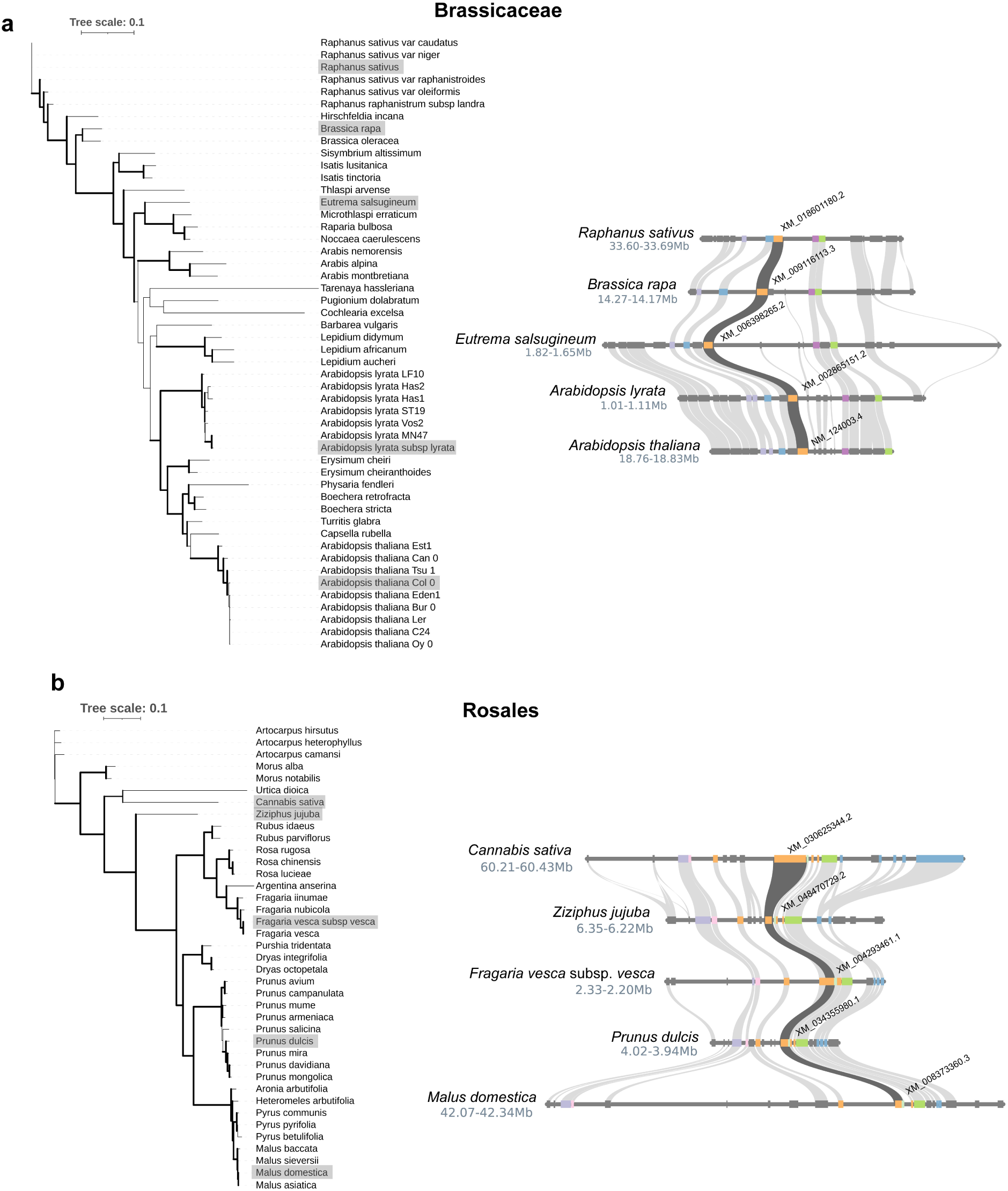
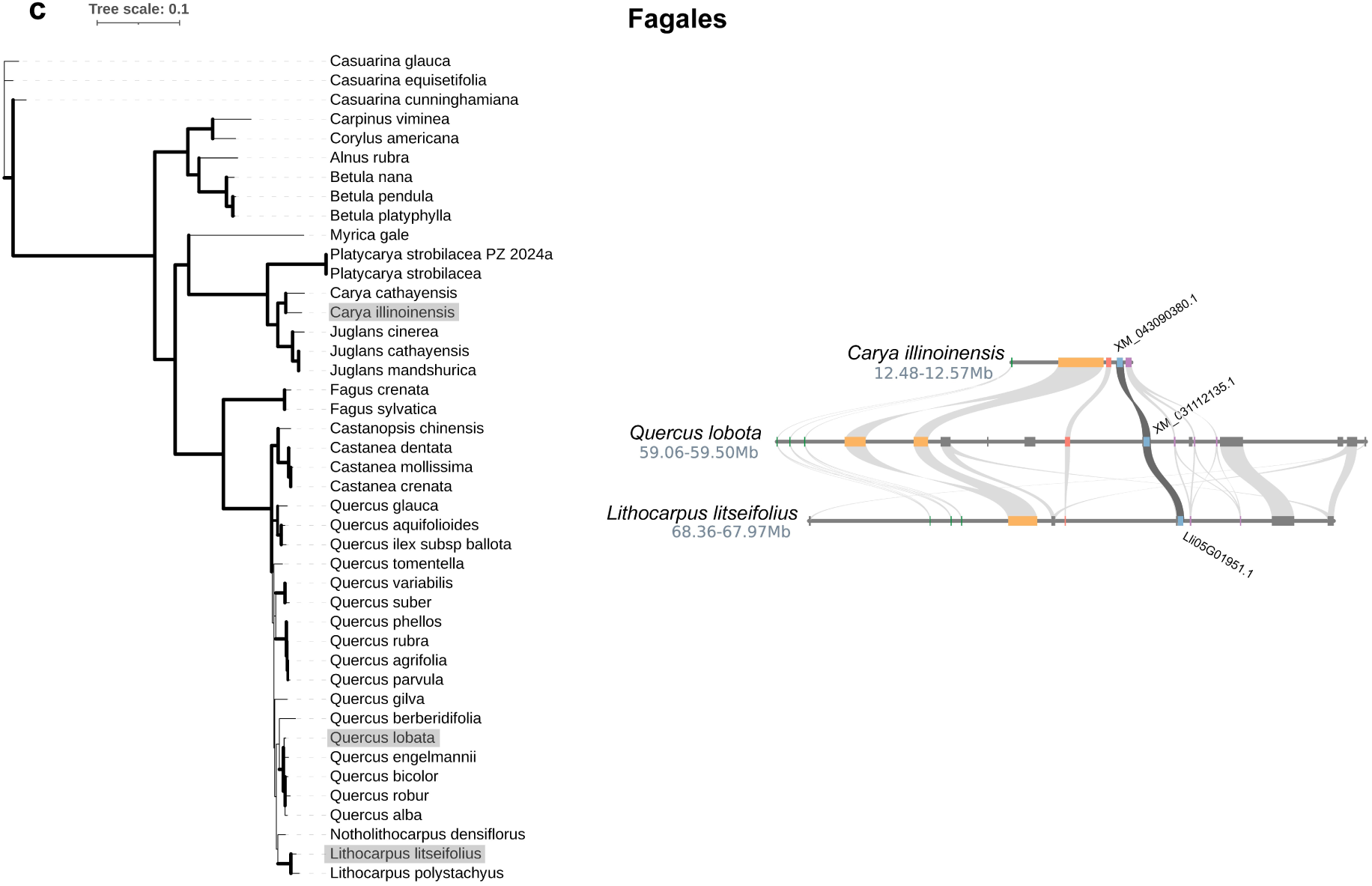
Phylogenetic and microsynteny analysis of FLS2 in Brassicaceae, Rosales and Fagales. **a** to **c,** Brassicaceae, Rosales and Fagales. **Left**: Maximum likelihood phylogeny constructed using single-copy, full-length FLS2 protein sequences from each phylogenetic group with 1000 bootstrap repeats. Branches supported by > 70% bootstrap values are indicated with thick dark lines. Species included in the microsynteny analysis are shaded gray. **Right**: Microsynteny analysis of FLS2 alongside ten upstream and downstream genes. Orthologous gene pairs are linked by light gray bands. FLS2 pairs are emphasized with dark gray bands, with NCBI identifiers labeled on the side. Only genomes with high-quality annotations are included in this analysis.

## References

1. Ngou, B. P. M., Ding, P. & Jones, J. D. G. Thirty years of resistance: Zig-zag through the plant immune system. Plant Cell 34, 1447–1478 (2022).

2. Ngou, B. P. M., Wyler, M., Schmid, M. W., Kadota, Y. & Shirasu, K. Evolutionary trajectory of pattern recognition receptors in plants. Nat Commun 15, 308 (2024).

3. Shiu, S.-H. & Bleecker, A. B. Receptor-like kinases from *Arabidopsis* form a monophyletic gene family related to animal receptor kinases. Proc Natl Acad Sci U S A 98, 10763–10768 (2001).

4. Albert, I. et al. An RLP23–SOBIR1–BAK1 complex mediates NLP-triggered immunity. Nat Plants 1, 1–9 (2015).

5. Yu, X., Feng, B., He, P. & Shan, L. From Chaos to Harmony: Responses and Signaling upon Microbial Pattern Recognition. Annual Review of Phytopathology 55, 109–137 (2017).

6. Chinchilla, D., Bauer, Z., Regenass, M., Boller, T. & Felix, G. The Arabidopsis Receptor Kinase FLS2 Binds flg22 and Determines the Specificity of Flagellin Perception. The Plant Cell 18, 465–476 (2006).

7. Matsui, S., et al. *Arabidopsis* SBT5.2 and SBT1.7 subtilases mediate C-terminal cleavage of flg22 epitope from bacterial flagellin. Nat Commun 15, 3762 (2024).

8. Sun, Y. et al. Structural Basis for flg22-Induced Activation of the Arabidopsis FLS2-BAK1 Immune Complex. Science 342, 624–628 (2013).

9. Trdá, L. et al. The grapevine flagellin receptor VvFLS2 differentially recognizes flagellin-derived epitopes from the endophytic growth-promoting bacterium *Burkholderia phytofirmans* and plant pathogenic bacteria. New Phytol 201, 1371–1384 (2014).

10. Trinh, J. et al. Variation in microbial feature perception in the Rutaceae family with immune receptor conservation in citrus. Plant Physiol 193, 689–707 (2023).

11. Felix, G., Duran, J. D., Volko, S. & Boller, T. Plants have a sensitive perception system for the most conserved domain of bacterial flagellin. The Plant Journal 18, 265–276 (1999).

12. Robatzek, S. et al. Molecular identification and characterization of the tomato flagellin receptor LeFLS2, an orthologue of *Arabidopsis* FLS2 exhibiting characteristically different perception specificities. Plant Mol Biol 64, 539–547 (2007).

13. Buscaill, P. & van der Hoorn, R. A. L. Defeated by the nines: nine extracellular strategies to avoid microbe-associated molecular patterns recognition in plants. Plant Cell 33, 2116–2130 (2021).

14. Colaianni, N. R. et al. A complex immune response to flagellin epitope variation in commensal communities. Cell Host Microbe 29, 635–649.e9 (2021).

15. Fürst, U. et al. Perception of *Agrobacterium tumefaciens* flagellin by FLS2XL confers resistance to crown gall disease. Nat. Plants 6, 22–27 (2020).

16. Wu, L., Xiao, H., Zhao, L. & Cheng, Q. CRISPR/Cas9-mediated generation of fls2 mutant in *Nicotiana benthamiana* for investigating the flagellin recognition spectrum of diverse FLS2 receptors. Plant Biotechnol J 20, 1853–1855 (2022).

17. Wei, Y. et al. An immune receptor complex evolved in soybean to perceive a polymorphic bacterial flagellin. Nat Commun 11, 3763 (2020).

18. Stevens, D. M. et al. Natural variation of immune epitopes reveals intrabacterial antagonism. Proc Natl Acad Sci U S A 121, e2319499121 (2024).

19. Tang, J. et al. Structural basis for recognition of an endogenous peptide by the plant receptor kinase PEPR1. Cell Res 25, 110–120 (2015).

20. Hebditch, M. & Warwicker, J. Charge and hydrophobicity are key features in sequence-trained machine learning models for predicting the biophysical properties of clinical-stage antibodies. PeerJ 7, e8199 (2019).

21. Osorio, D., Rondón-Villarreal, P. & Torres, R. Peptides: A Package for Data Mining of Antimicrobial Peptides. The R Journal 7, 4 (2015).

22. Homma, F., Huang, J. & van der Hoorn, R. A. L. AlphaFold-Multimer predicts cross-kingdom interactions at the plant-pathogen interface. Nat Commun 14, 6040 (2023).

23. Fawcett, T. An introduction to ROC analysis. Pattern Recognition Letters 27, 861–874 (2006).

24. Torres Ascurra, Y. C., et al. Functional diversification of a wild potato immune receptor at its center of origin. Science 381, 891–897 (2023).

25. Prigozhin, D. M. & Krasileva, K. V. Analysis of intraspecies diversity reveals a subset of highly variable plant immune receptors and predicts their binding sites. The Plant Cell 33, 998–1015 (2021).

26. Kuang, H., Woo, S.-S., Meyers, B. C., Nevo, E. & Michelmore, R. W. Multiple Genetic Processes Result in Heterogeneous Rates of Evolution within the Major Cluster Disease Resistance Genes in Lettuce[W]. The Plant Cell 16, 2870–2894 (2004).

27. Gong, Z. & Han, G.-Z. Flourishing in water: the early evolution and diversification of plant receptor-like kinases. Plant J 106, 174–184 (2021).

28. Rentel, M. C., Leonelli, L., Dahlbeck, D., Zhao, B. & Staskawicz, B. J. Recognition of the Hyaloperonospora parasitica effector ATR13 triggers resistance against oomycete, bacterial, and viral pathogens. Proceedings of the National Academy of Sciences 105, 1091–1096 (2008).

29. Thomas, N. C. et al. The rice XA21 ectodomain fused to the *Arabidopsis* EFR cytoplasmic domain confers resistance to *Xanthomonas oryzae pv. oryzae*. PeerJ 6, e4456 (2018).

30. Wulff, B. B. H., Thomas, C. M., Smoker, M., Grant, M. & Jones, J. D. G. Domain Swapping and Gene Shuffling Identify Sequences Required for Induction of an Avr-Dependent Hypersensitive Response by the Tomato Cf-4 and Cf-9 Proteins. Plant Cell 13, 255–272 (2001).

31. Mueller, K. et al. Chimeric FLS2 Receptors Reveal the Basis for Differential Flagellin Perception in Arabidopsis and Tomato. The Plant Cell 24, 2213–2224 (2012).

32. Helft, L., Thompson, M. & Bent, A. F. Directed Evolution of FLS2 towards Novel Flagellin Peptide Recognition. PLOS ONE 11, e0157155 (2016).

33. Seong, K. et al. Engineering the plant intracellular immune receptor Sr50 to restore recognition of the AvrSr50 escape mutant. 2024.08.07.607039 Preprint at 10.1101/2024.08.07.607039 (2024).

34. Cruz, D. G. D. L., Zdrzałek, R., Banfield, M. J., Talbot, N. J. & Moscou, M. J. Molecular mimicry of a pathogen virulence target by a plant immune receptor. bioRxiv 2024.07.26.605320 (2024) doi:10.1101/2024.07.26.605320.

35. Snoeck, S. et al. Leveraging coevolutionary insights and AI-based structural modeling to unravel receptor–peptide ligand-binding mechanisms. Proceedings of the National Academy of Sciences 121, e2400862121 (2024).

36. Kourelis, J., Marchal, C., Posbeyikian, A., Harant, A. & Kamoun, S. NLR immune receptor– nanobody fusions confer plant disease resistance. Science 379, 934–939 (2023).

37. Tamborski, J., Seong, K., Liu, F., Staskawicz, B. J. & Krasileva, K. V. Altering Specificity and Autoactivity of Plant Immune Receptors Sr33 and Sr50 Via a Rational Engineering Approach. MPMI 36, 434–446 (2023).

38. Zhang, X. et al. The synthetic NLR RGA5HMA5 requires multiple interfaces within and outside the integrated domain for effector recognition. Nat Commun 15, 1104 (2024).

39. Katoh, K. & Standley, D. M. MAFFT Multiple Sequence Alignment Software Version 7: Improvements in Performance and Usability. Molecular Biology and Evolution 30, 772–780 (2013).

40. Minh, B. Q. et al. IQ-TREE 2: New Models and Efficient Methods for Phylogenetic Inference in the Genomic Era. Molecular Biology and Evolution 37, 1530–1534 (2020).

41. Guindon, S. et al. New Algorithms and Methods to Estimate Maximum-Likelihood Phylogenies: Assessing the Performance of PhyML 3.0. Systematic Biology 59, 307–321 (2010).

42. Letunic, I. & Bork, P. Interactive Tree of Life (iTOL) v6: recent updates to the phylogenetic tree display and annotation tool. Nucleic Acids Research 52, W78–W82 (2024).

43. Yang, Z., Nielsen, R., Goldman, N. & Pedersen, A.-M. K. Codon-Substitution Models for Heterogeneous Selection Pressure at Amino Acid Sites. Genetics 155, 431–449 (2000).

44. Huerta-Cepas, J., Serra, F. & Bork, P. ETE 3: Reconstruction, Analysis, and Visualization of Phylogenomic Data. Mol Biol Evol 33, 1635–1638 (2016).

45. Tang, H., et al. JCVI: A versatile toolkit for comparative genomics analysis. iMeta 3, e211 (2024).

46. Helft, L. et al. LRR Conservation Mapping to Predict Functional Sites within Protein Leucine-Rich Repeat Domains. PLOS ONE 6, e21614 (2011).

47. Gu, Z., Eils, R. & Schlesner, M. Complex heatmaps reveal patterns and correlations in multidimensional genomic data. Bioinformatics 32, 2847–2849 (2016).

48. Mirdita, M. et al. ColabFold: making protein folding accessible to all. Nat Methods 19, 679– 682 (2022).

49. Abramson, J. et al. Accurate structure prediction of biomolecular interactions with AlphaFold 3. Nature 630, 493–500 (2024).

